# *Portiera* gives new clues on the evolutionary history of whiteflies

**DOI:** 10.1101/2020.06.17.158493

**Authors:** D. Santos-Garcia, N. Mestre-Rincon, D. Ouvrard, E. Zchori-Fein, S. Morin

## Abstract

Whiteflies (Hemiptera: Sternorrhyncha: Aleyrodidae) are a superfamily of small phloem-feeding insects. Their taxonomy is currently based on the morphology of nymphal stages that display phenotypic plasticity, which produces inconsistencies. To overcome this limitation, we developed a new phylogenetic framework that targets five genes of *Candidatus* Portiera aleyrodidarum, the primary endosymbiont of whiteflies. *Portiera* lineages have been co-diverging with whiteflies since their origin and therefore reflect their host evolutionary history. We also studied the origin of stability and instability in *Portiera* genomes by testing for the presence of two alternative gene rearrangements and the loss of a functional polymerase proofreading subunit (*dnaQ*), previously associated with genome instability. We present two phylogenetic reconstructions. One using the sequences of all five target genes from 22 whitefly species belonging to 17 genera. The second uses only two genes to include additional published *Portiera* sequences of 21 whitefly species, increasing our sampling size to 42 species from 25 genera. The developed framework showed low signal saturation, specificity to whitefly samples, and efficiency in solving inter-genera relationships and standing inconsistencies in the current taxonomy of the superfamily. Genome instability was found to be present only in the Aleurolobini tribe containing the *Singhiella, Aleurolobus* and *Bemisia* genera. This suggests that *Portiera* genome instability likely arose in the Aleurolobini tribe’s common ancestor, around 70 Mya. We propose a link between the switch from multi-bacteriocyte to a single-bacteriocyte mode of inheritance in the Aleurolobini tribe and the appearance of genome instability in *Portiera*.

## 1 Introduction

The order Hemiptera is the largest monophyletic group of hemimetabolous (without metamorphosis) insects and is, characterized by specialized piercing-sucking mouth-parts (“rostrum”) (Grimaldi and Engel, 2005). The hemipteran order is divided into four suborders: Sternorrhyncha, Auchenorryncha, Heteroptera, and Coleorrhyncha. Most hemipteran insects feed on plant sap (phloem and xylem), but some heteropteran insects are predatory (Grimaldi and Engel, 2005; Cryan and Urban, 2012). The Sternorrhyncha suborder is usually divided into the Aphidinea lineage containing the Aphidoidea (aphids) and Coccoidea (scale insects) superfamilies, and the Psyllinea lineage containing the Psylloidea (psyllids) and Aleyrodoidea (whiteflies) superfamilies (Shcherbakov, 2000).

The Aleyrodoidea superfamily is composed of one family, the Aleyrodidae, which is divided into two extant subfamilies, the Aleurodicinae and the Aleyrodinae, and an extinct one, the Bernaeinae. The Aleurodicinae subfamily is composed of 20 genera, mainly distributed in Neotropical/Australasian regions (Charles, 2010). The Aleyrodinae, with 140 described genera, is a more diverse and globally distributed (Manzari and Quicke, 2006). It includes the major pest species *Bemisia tabaci* and *Trialeurodes vaporariorum*. Although the extant whitefly subfamilies were reported to have their origin in the Middle Cretaceous (Campbell *et al*., 1994), the first fossils of the present extant families were dated to the Lower Cretaceous (Drohojowska and Szwedo, 2015). During that period, whiteflies were associated with gymnosperm forests and/or pro-angiosperms, in contrast to extant whitefly species, which feed mainly on angiosperms. It is assumed that the emergence of angiosperms in the Lower Cretaceous has promoted the radiation of the whitefly superfamily due to the opening of new environmental niches. This event triggered their diversification and speciation along with the angiosperms (Middle-Upper Cretaceous), leading to the emergence of the modern whitefly species (Drohojowska and Szwedo, 2015).

Whiteflies are the less diverged superfamily among the Sternorrhyncha. Their taxonomy is based mainly on the fourth instar nymph morphology, which is quiescent and known as “puparium”, instead of the more commonly used adult phase (Martin and Mound, 2007; Evans, 2007). The “puparium” stage presents phenotypic plasticity with phenotypes affected both by the identity of the host plants and different abiotic factors (Manzari and Quicke, 2006; Martin and Mound, 2007). Also, most of the phylogenetic research so far has focused on relatively few species with significant economic impacts, such as the *B. tabaci* species complex (Brown *et al*., 1995; Frohlich *et al*., 1999; Boykin *et al*., 2007; De Barro *et al*., 2011; Lee *et al*., 2013; Hsieh *et al*., 2014; Wang *et al*., 2014). To the best of our knowledge, only four published studies have tried to establish broad phylogenetic relationships among the different whitefly genera. Three of them were based on molecular phylogenetics (Thao and Baumann, 2004a; Ovalle *et al*., 2014; Dickey *et al*., 2015) while the third utilized morphological cladistics (Manzari and Quicke, 2006). Thao and Baumann (2004a) used *Portiera*’s *16S* and *23S* rRNA genes to classify 20 species from 12 genera and reported some incongruence between taxonomical names and molecular data. Both Ovalle *et al.* (2014) and Dickey *et al.* (2015) works used the 5’ region of the mitochondrial cytochrome oxidase 1 (*mtCOI*) barcode to classify 20 species from 9 genera and 104 species from 24 genera, respectively. However, both studies showed low support at inner nodes and highlighted the limited capacity of the *mtCOI* barcode to establish inter-genera relationships. More recently, Manzari and Quicke (2006) used 94 morphological characters present in 439 whitefly species (from seven Aleurodicinae and 117 Aleyrodinae genera) to reconstruct the species phylogenetic relationships. The authors conclude that in many cases, the ability of the “puparium” morphological characters to provide species definition is likely to be limited and might not even be genera specific. Therefore, it is clear that the classification and diversity of whitefly superfamily need a reassessment.

Most sternorrhynchan insects, including whiteflies, present obligatory bacterial symbionts harbored within specialized cells, termed bacteriocyte. These primary intracellular symbionts, commonly named P-endosymbionts, generally complement their hosts’ restricted diets (plant sap) (Hansen and Moran, 2014). P-endosymbionts exhibit mother-to-offspring vertical transmission, which promotes their co-speciation with their hosts and enable them to reflect their hosts’ phylogenetic history (co-cladogenesis) (De Vienne *et al*., 2013). The genome of P-endosymbionts has been reduced to a basic set of genes responsible for maintaining the symbiotic relationship and minimal cell machinery. Other characteristics include the absence (nearly) of gene duplications, mobile elements, and acquisition of new genetic material (no gene flow) (Latorre and Manzano-Marín, 2016). These characteristics make the genomes of P-endosymbionts a valuable resource for reconstructing their hosts phylogenetic relationships, as already have been demonstrated in aphids (Martinez-Torres *et al*., 2001; Jousselin *et al*., 2009; Nováková *et al*., 2013; Meseguer *et al*., 2015, 2017) and psyllids (Hall *et al*., 2016).

The P-endosymbiont of whiteflies is *Candidatus* Portiera aleyrodidarum (hereafter *Portiera*) (Thao and Baumann, 2004b). *Portiera* forms a monophyletic clade together with *Ca*. Carsonella ruddii, the P-endosymbiont of psyllids. Based on molecular data, it has been proposed that the ancestral symbiosis was established in the Psyllinea lineage, before their divergence into the Aleyrodoidea and Psylloidea lineages (Santos-Garcia *et al*., 2014). Therefore, *Portiera* has been co-diverging with their whiteflies hosts since their origin, reflecting this way both the hosts’ phylogenetic relationships and divergence time (Santos-Garcia *et al*., 2015). Until now, only three *Portiera* genomes from species others than *B. tabaci* have been sequenced. As other P-endosymbionts, these three *Portiera* have maintained a genome stasis for more than 135 million years (Myr) (Sloan and Moran,2013; Santos-Garcia *et al*., 2015). In contrast, *Portiera* genomes from the *B. tabaci* species complex, although syntenic among themselves, have lost the synteny with the other three published *Portiera* genomes. The genome rearrangements of *Portiera* from *B. tabaci* is correlated with a massive loss of genes required for correct DNA replication and the repair machinery. These losses include the DNA polymerase III subunit epsilon *dnaQ*, which is required for repairing spontaneous mutations (proofreading activity) (Sloan and Moran, 2013; Santos-Garcia *et al*., 2015). Currently, it is not clear if the above mentioned genome instability is a unique characteristic of *Portiera* from *B. tabaci* or a more general phenomenon present in other lineages.

In this work, we explored the use of *Portiera* as a tool for determining relationships among whiteflies. First, we used up to five *Portiera* genes to reconstruct the phylogeny and divergence of 42 whitefly species belonging to 25 different genera. Freshly collected, ethanol stored, and museum *exsiccata* samples were used to ensure our framework’s robustness. Our analysis indicated that phylogenetics based on *Portiera* gene sequences is a promising approach as it shows low signal saturation and can efficiently solve inter-genera relationships. Second, we used genome sequencing and PCR amplification to screen for genome rearrangements and the presence/absence of a functional dnaQ gene in *Portiera* to understand better the origin of the genome instability found in some *Portiera* lineages. Our analysis indicated that *Portiera* genome instability originated once in the Aleurolibini tribe ancestor. The possible link between a newly emerged mode of bacteriocyte transmission in the Aleurolibini tribe and the appearance of genome instability is discussed.

## 2 Material and Methods

### Whiteflies collections and gDNA extraction

A total of 29 samples, accounting for 25 different whitefly species, were obtained. Whitefly samples were from different sources: freshly collected and stored in ethanol until use (adult insects), Prof. Dan Gerling’s collection (adult insects conserved in ethanol), and *exsiccate* collection samples from the Natural History Museum (NHM) in London (nymphs were removed from dry leaves and sent in ethanol). Before any genomic DNA extraction was performed, five adult insects (or nymphs from the NHM collection) were rehydrated by consecutive passes in 70%, 50%, 30%, 0% v/v ethanol solutions in sterile water. Whiteflies were transferred to a new 1.5 ml tube containing 80*µ*l lysis buffer T1 and were homogenized with 1.4 mm zirconia beads (CK14, Bertin Instruments) using a bead-beater (Minilys, Bertin Instruments). Genomic DNA (gDNA) was extracted with NucleoSpin Tissue XS (Macherey-Nagel) following the manufacturer instructions. For the NHM samples, a nondestructive method was used whenever possible. Nymphs were incubated overnight (56°C) in 80*µ*l lysis buffer T1 and 8*µ*l Proteinase K (20*µ*g/*µ*l). gDNA was extracted from the lysis buffer using the NucleoSpin Tissue XS standard protocol. The nymphs were recovered, cleaned with sterile water, and stored in fresh ethanol. gDNA from seven samples that had less than five individuals were subjected to whole-genome amplification (GenomiPhi V2, GE Healthcare), following manufacturer instructions, to ensure sufficient material. For Illumina sequencing, *Singhiella simplex* adults were accidentally collected together with *Pealius mori* adult whiteflies in July 2018 from *Ficus benjamina* (GPS coordinates 31.904511;34.804562) and stored in ethanol. Later on, whiteflies were rehydrated and sexed. Female abdomens (50) were dissected under a stereo-microscope using autoclaved 1X Phosphate-buffered saline. Abdomens were homogenized with a bead-beater, and gDNA was extracted with NucleoSpin Tissue XS, as described above. Whole-genome shotgun sequencing was performed in a NovSeq 6000u sing a TruSeq DNA PCR Free Library (2×150bp) at Macrogen Europe.

### PCR screening, sequencing, and ancestral estate reconstruction

For the PCR screening, five genes present in all insect endosymbionts showing extremely reduced genomes were selected (Moran and Bennett, 2014). These genes are widely used as bacterial phylogenetic markers and includes the *16S* and *23S* ribosomal RNAs (rRNAs), the chaperonins *groEL* and *dnaK*, and the RNA polymerase sigma factor *rpoD*. We manually designed *Portiera* specific universal primers using available *Portiera* genomes from both the Aleyrodinae (*B. tabaci* and *T. vaporariorum*) and Aleurodicinae (*Aleurodicus dispersus* and *A. floccissimus*) subfamilies in UGENE v1.28.1 (Okonechnikov *et al*., 2012) (Table S1). Primers melting temperature (Tm), off-targets, and possible primer-dimer interactions were computed with Primer3 software implemented on https://eu.idtdna.com/calc/analyzer.

Primers (0.5 mM each) were mixed with the KAPA2G Robust HotStart ReadyMix (Kapa Biosystems) inside a DNA/RNA UV-Cleaner cabinet (UVC/T-AR). PCR was performed using the following general profile: 95°C for 5 min,[95°C for 30 sec, Tm°C for 15 sec, 72°C for 1 min]x35, 72°C for 5 min. Annealing temperature (Tm) was set up for each primer set according to Primer3 predictions (Table S1). When required, the temperature was adjusted trying 5°C above or below of the predicted Tm.

PCR product size was confirmed by electrophoresis using 1% agarose gel, purified with DNA Clean & Concentrator 5 (Zymo Research), and sequenced by Sanger technology in both directions at Macrogen Europe. For each amplicon, sequences were quality screening/clipping and their consensus alignment performed with the Staden Package (Bonfield and Whitwham, 2010).

In parallel, we designed primers that target the DNA polymerase III subunit epsilon *dnaQ*. Also, we targeted two regions with different gene order in *Portiera* of *B. tabaci, lepA*-*groEL* (A_*Bt*_) and*secA*-*leuC* (B_*Bt*_), compared to the ancestral gene order found in other sequenced *Portiera, groEL*-*rpsA* (A) and *leuC* -*leuD* (B). Primer design and PCR reactions were conducted as described above using the predicted Tm (Table S1). PCR products were visualized by electrophoresis using a 1%. Some obtained amplicons were Sanger sequenced to validate the amplified region.

Additionally, for each whitefly species collected, the 5’ region of the mitochondrial cytochrome oxidase 1 (*mtCOI*) gene was amplified using the universal primers LCO1490 (F) and HCO2198 (R) (Folmer *et al*., 1994). In cases were this set of primers failed to amplify, the C1-J-2195 (F) and L2-N-3014 (R) primer set targeting the 3’ region was used (Frohlich *et al*., 1999). In species were both sets of primers failed to amplify and *mtCOI* sequences were available at public databases, species-specific sets were designed (Table S1). PCR conditions and procedures were the same as described above.

### *Portiera* lineages Divergence Dating

Two datasets were used to infer the divergence time of the whitefly species analyzed. The first dataset incorporated sequences of *Portiera 16S* and *23S* rRNA genes amplified in this study, *16S* and *23S* rRNA gene sequences generated by Thao and Baumann (2004b), and *16S* and *23S* rRNA gene sequences extracted from the downloaded transcriptomes/genomes. The final dataset contained 59 sequences from 45 different species (including six belonging to the *B. tabaci* species complex). The second dataset integrated the sequences of the *16S* and *23S* rRNA genes with those of the three coding genes: *dnaK, rpoD*, and *groEL*. It contained 32 sequences from 29 whitefly species, mostly obtained in this study plus few that were acquired from public transcriptomes/genomes. In both datasets analysis, orthologous genes extracted from *Chromohalobacter salexigens* DSM3043 (NC 007963.1) were used as outgroups.

The *16S* and *23S* rRNA genes were aligned with R-Coffee v11.00.8cbe486 (-mode=rmcoffee -iterate=100) (Notredame *et al*., 2000) and pruned with Gblocks v0.91b allowing half of gap positions (-t=d -b5=h) (Castresana, 2000). The three coding genes (*dnaK, rpoD*, and *groEL*) were codon-aligned with MACSE v2.03 (-prog alignSequences -gc def 11) (Ranwez *et al*., 2018) and pruned with Gblocks v0.91b (-t=c -b5=h). The 19 obtained *mtCOI* gene sequences (5’ region) were aligned in the same way but using the invertebrate mitochondrial code in MACSE v2.03 and no gaps allowed in Gblocks v0.91b. Substitution saturation was assessed using the pruned alignments as an input for Xia’s test implemented in DAMBE v7.2.3 (Xia, 2018) (executed under wine v1.6.2-0ubuntu14.2).

BEAST v2.5.2 (Bouckaert *et al*., 2014) was used to infer a Bayesian posterior consensus tree and the divergence time of the different nodes as previously described (Santos-Garcia *et al*., 2015). Detailed procedures of divergence dating can be found at Supplementary Material and Methods.

### Whole Genome Shotgun sequencing, genome assembly and annotation of the Singhiella simplex and Pealius mori joint sample

NovaSeq sequencing produced 75,274,888 raw reads that were quality screened with Trimmomatic v0.33 (same parameters as described above). Possible polyGs produced by the NovaSeq platform were trimmed with fastp v0.19.7 (Chen *et al*., 2018). Cleaned reads were classified with Kraken v2.0.6-beta and the custom database described above. All reads assigned to *Portiera, Halomonadaceae*, or *Oceanospirillales* were extracted and assembled with SPAdes v3.13.0 (–sc –careful) (Bankevich *et al*., 2012). Three contigs larger than 60Kb (385Kb in total) and ∼100x coverage plus several contigs between 80Kb and 5Kb (420Kb in total) and ∼600X coverage were recovered. Kraken2 classification and coverage suggested two putative *Portiera* populations. To screen for possible a *Portiera* other than that of *Singhiella simplex*, all sequences obtained during the PCR screening were used as a query in a BLASTN search against the obtained contigs. BLASTN results confirmed that two different *Portiera* genomes were present. Some large contigs with ∼100x coverage had perfect match to the *Portiera* amplified genes from *Pealius mori*. Several smaller contigs with a coverage of ∼600X had perfect match to the amplified *Portiera* genes from *S. simplex*. This confirmed that some *P. mori* individuals were collected together with *S. simplex*, probably due to the ability of both whitefly species to exploit *Ficus benjamina* as a host-tree.

As a result, the Kraken2 database was rebuilt including the obtained contigs, and cleaned reads were re-classified. *Portiera* reads were re-assembled separately according to their whitefly host with SPAdes v3.13.0. SSPACE v3 (Boetzer *et al*., 2011) and GapFiller v1.10 (Boetzer and Pirovano, 2012) were used for scaffolding and gap-filling of the re-classified reads. Gap5 from the Staden package was used to evaluate the assemblies quality, to detect the presence of chimeras and miss-assemblies, to join contings manually (when possible), and to check for circular contigs. The first genome to be assembled was that of *Portiera* from *P. mori*. It produced a closed circular contig without requiring any iterative mapping step. In contrast, the *Portiera* genome from *S. simplex* remained as 9 contigs after several rounds of iterative mapping, discarding at each round every sequence (if present) with a significant (90% identity threshold) match to the *Portiera* genome of *P. mori*. In brief, iterative mapping was run as follows: cleaned reads were mapped against the assembled contigs of *S. simplex* with Bowtie v2.3.5.1 (Langmead and Salzberg, 2012). Reads without a minimum overlap of 50% and 90% identity to the contigs were discarded. Surviving reads were added to the pool of putative *Portiera* reads from *S. simplex*. The reads were mapped to the contigs with MIRA v4.9.6 (Chevreux, B., Wetter, T. and Suhai,1999), and then imported to Gap5 for manual joining/gap closure. Both final assemblies were corrected with Pilon v1.23 (–fix all,amb) (Walker *et al*., 2014) and the clean classified reads. Finally, the annotation of the genomes was performed with prokka v1.14.5 (Seemann, 2014), using all available *Portiera* genomes for building the protein database of primary annotation (–protein). Obtained annotations were manually inspected and curated in Artemis v1.5 (Rutherford *et al*., 2000).

### *Portiera* lineages Comparative Genomics

Proteomes of *Portiera* from *S. simplex* (ERZ1272841), *P. mori* (ERZ1272840), *B. tabaci* species - MEAM1 (NC 018677.1), MED (NC 018676.1), and Asia II 3 (NZ CP016327.1), *T. vaporariorum* (LN649236.1), *A. dispersus* (LN649255.1) and *A. floccissimus* (LN734649.1) were extracted with a custom python script. Orthologous clusters of proteins (OCPs) were calculated with OrthoFinder v2.3.3 (-M msa -S mmseqs -T iqtree) (Emms and Kelly, 2019). Obtained OCPs were manually curated based on protein annotations. Shared and specific OCPs were plotted with UpsetR Conway *et al*. (2017). Synteny between *Portiera* genomes, based on 230 single-copy core OCPs (from 235), was plotted with genoPlotR (Guy *et al*., 2010). Finally, metabolic potential comparisons were performed with Pathway Tools v23.5 Karp *et al.* (2002).

Curated OCPs were converted into a binary matrix (presence/absence) and species-specific OCPs annotated as hypothetical proteins were discarded (21 OCPs). The binary matrix and the species tree obtained with OrthoFinder v2.3.3 were used as inputs for COUNT v10.04 (Csuos, 2010) to reconstruct the gene losses history during *Portiera* evolution. The reconstruction was performed under a posterior algorithm and allowed only gene losses.

Results from the *dnaQ* screening, codified as a binary matrix, were used together with the Bayesian trees obtained (the tree based only on two rRNA genes and the tree based on the five gene set) as an input for the Ancestral Character Estimation (ace) function implemented in ape (R package) (R Core Team, 2018; Paradis and Schliep, 2019). The presence of *dnaQ* on the internal nodes from both datasets was estimated using a maximum likelihood (ML) approach as a discrete character and a model assuming only gene losses. Phylogenetic trees with *dnaQ* presence probabilities were plotted with ape.

### *Portiera* lineages Molecular Evolution

Synonymous (*dS*) and non-synonymous (*dN*) substitution ratios, nucleotide substitutions per site per year (*dS/t* and *dN/t*), and omega (*ω*) values were calculated as previously described in Santos-Garcia *et al*. (2015) using Codeml from PAML v4.7 package (Yang, 2007). The full procedure can be found at Supplementary Material and Methods.

### Repeats and intergenic regions analysis

Repeats annotation and intergenic regions comparative analysis can be found at Supplementary Material and Methods.

### Transcriptome and Mitochondrion assembly, data retrieval, and mitochondrial molecular evolution

*Dialeurodes citri* and *B. tabaci* SSA1 transcriptome assembly procedures, mitochondrial genomes assembly and annotation of *S. simplex* and *P. mori*, and *Portiera* genomes and whiteflies’ mitogenomes retrieval can be found at Supplementary Material and Methods.

## 3 Results

### *Portiera* marker genes amplification

gDNA was extracted from 26 of the 29 collected samples, standing for 22 whitefly species from 17 genera (Table S1). We failed to obtain gDNA from three NHM collection exsiccate samples (Table S1), even when applying a whole genome amplification approach. The five sets of primers that target *Portiera* genes successfully amplified in 25 samples. One sample, *Bemisia reyesi* JHM 7496, was excluded from further analysis because we could not amplify the target regions of the *23S* rRNA and *rpoD* genes. The primers sets targeting the *mtCOI* gene had lower efficiency and produced the expected PCR fragments in 18 of the 26 samples. The LCO1490/HCO2198 set produced a PCR fragment in 15 samples but two of them were parasitoid wasp sequences instead of whitefly sequences. The C1-J-2195/L2-N-3014 produced a PCR fragment in one sample. The other four successful amplifications used species-specific *mtCOI* primers. The five amplified *Portiera* genes did not show signatures of substitution saturation in their phylogenetic signal, and presented saturation values that were lower than those calculated for the full set of mitochondrial genes including the *mtCOI* amplicon (Table S3). In fact, the third codon positions of the *mtCOI* full gene and amplicon are completely saturated.

### Establishing *Portiera* phylogenetic relationships and divergence dating

The number of taxa and genes that are used in phylogenetic analysis can affect their outcome (Nabhan and Sarkar, 2012). For that reason, we conducted two phylogenetic reconstructions. One that used the sequences of all five target genes in *Portiera*: *16S* and *23S* rRNA, *dnaK, rpoD*, and *groEL*. The second analysis used only the two rRNA genes. This allowed us to add sequences of 20 whitefly species from 13 genera that were reported by Thao and Baumann (2004a).

In both phylogenetic trees, the Aleyrodinae subfamily outcompeted the Aleurodicinae subfamily in the number of analyzed species. The Aleurodicinae was represented by species from the genera *Paraleyrodes* and *Aleurodicus*. In both trees, the Aleyrodinae species clustered by genus, four major clusters could be identified and similar clustering pattern was obtained at the genera level (Figure 1 and 2). Some variation between the trees was observed in the orange cluster. In the tree that was based on five *Portiera* genes, the orange cluster was found to be the most basal branch and only contained one species, *T. vaporariorum* (Figure 1). In the tree that was based only on the two rRNA genes, the orange cluster (containing this time five species) was integrated within the purple cluster, and was close to the *Aleyrodes* clade (Figure 2). These topological inconsistencies likely result from the different taxon sampling in the two analyses. The tree that was based on five *Portiera* genes was well supported and most of the nodes presented posterior values greater than 0.9 (Figure 1). In contrast, the tree that was based on two rRNA genes had a large number of nodes with posterior values below 0.8, especially at some inner branches (Figure 1). Some of the low posterior support values were associated with potential species complexes: *tabaci, Aleyrodes singularis*/*proletella* and *Neomaskiella andropogonis*. Noticeable, some inconsistencies in taxonomy were also present in both trees. For example, *Aleuroviggianus adanaensis* was almost identical to *Tetraleurodes bicolor* at the sequence level and some species from the genera *Tetraleurodes, Trialeurodes*, and *Dialeurodes* were distributed among different clades.

**Figure 1:**
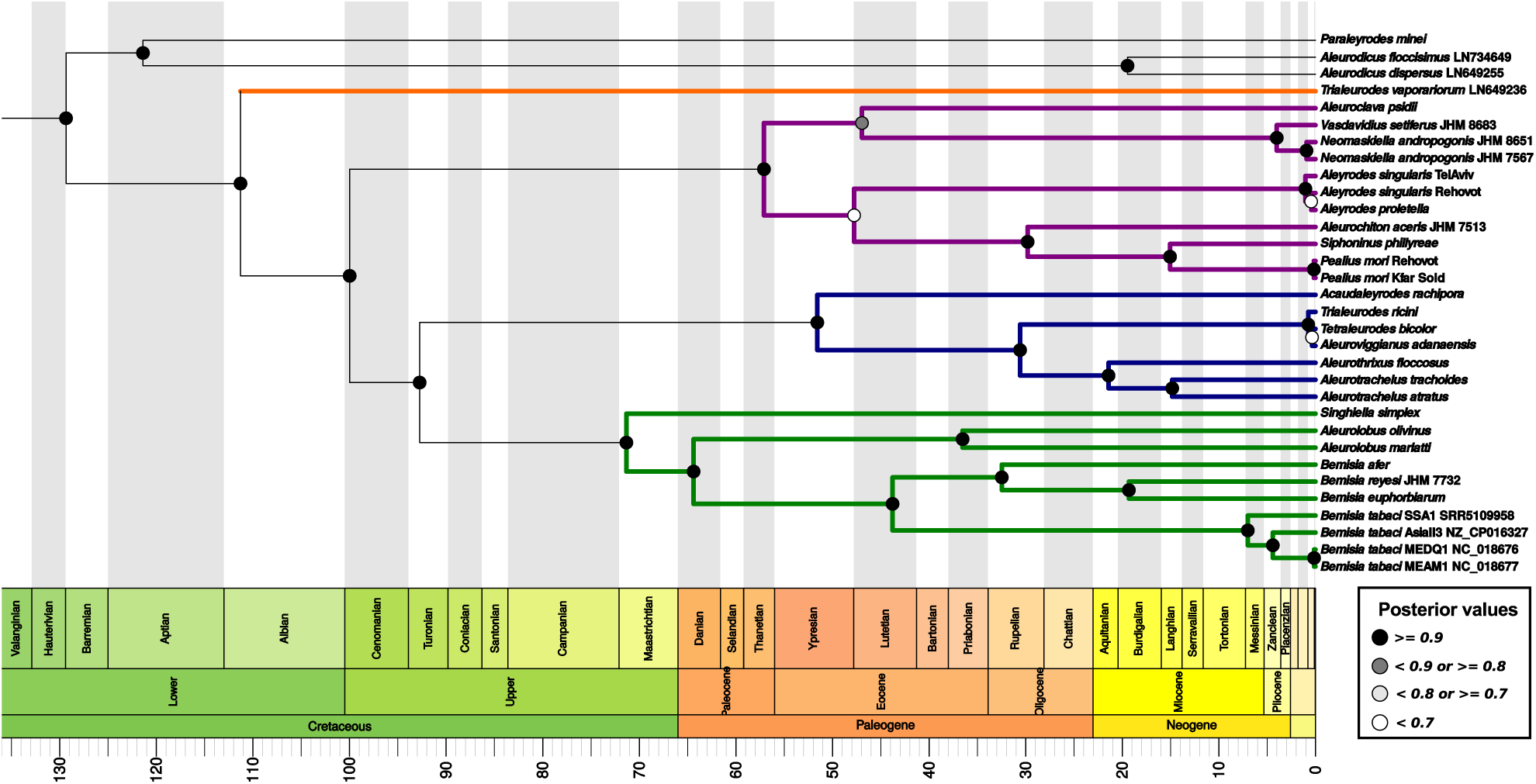
BEAST2 inferred *Portiera* chronogram based on two rRNA (*16S* and *23S*) and three coding genes (*groEL, rpoD*, and *dnaK*). Colored branches highlight the four major clades in the Aleyrodinae subfamily. Branch lengths are displayed in Million years. Period, Epoch, and Age are according to the geological time scale standards. *Chromohalobacter salexigens* DSM3043 was used as outgroup but is not displayed for plotting reasons.

**Figure 2:**
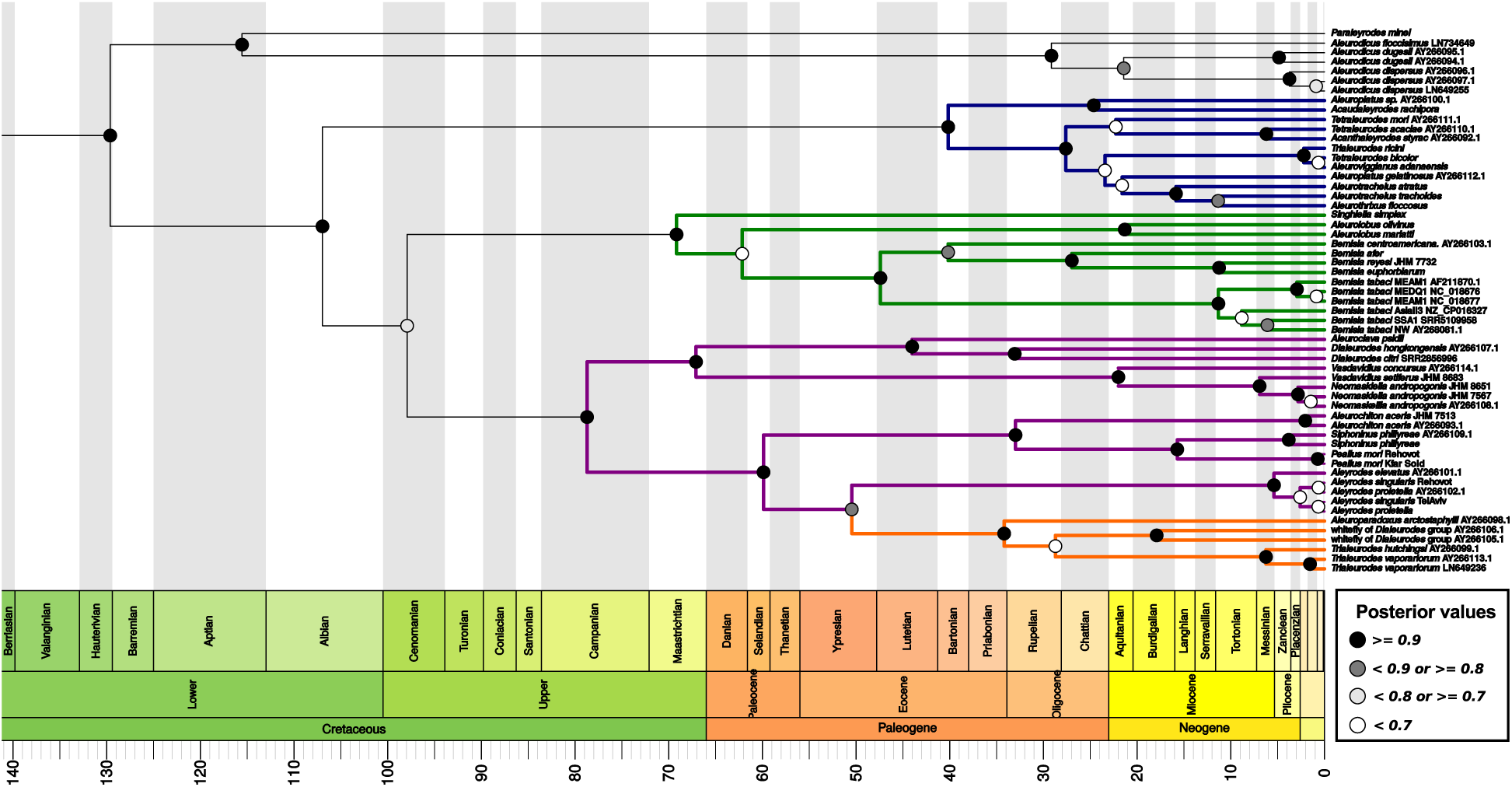
BEAST2 inferred *Portiera* chronogram based on two rRNA genes (*16S* and *23S*). The sequences were generated in this work and in Thao and Baumann (2004b). Colored branches highlight the four major clades in the Aleyrodinae subfamily Branch lengths are displayed in Million years. Period, Epoch, and Age are according to the geological time scale standards. *Chromohalobacter salexigens* DSM3043 was used as outgroup but is not displayed for plotting reasons.

Among the Aleurodicinae subfamily, *Paraleyrodes minei* was the first species to diverge, around 119.68 Mya (Million years ago) (102.74-133.42 95%HPD) or 112.6 Mya (85.49-133.31 95%HPD) according to the five genes- or two genes-based trees, respectively (Figure 1 and 2). The divergence of *Aleurodicus dispersus* from *Aleurodicus floccissimus* was estimated to be around 20.35 Mya (9.35-33.28 95%HPD) and 30.21 Mya (15.02-47.18 95%HPD) for the five genes- and two genes-based trees, respectively (Figure 1 and 2). These dates were in agreement with previous estimates Santos-Garcia *et al.* (2015). In the Aleyrodinae subfamily, despite the topological differences between the two trees, the estimated time of the first cladogenetic event (the first splitting after divergence from the main branch) was similar for the blue, green and purple clusters. The estimated divergence dates for the orange cluster were not comparable between the two datasets. However, if we consider the split between the green and purple/orange clusters in the two rRNA genes-based tree as the origin of the lineage leading to *T. vaporariorum*, then, the estimation for *T. vaporariorum* are quite similar: 97.36 Mya (76.14-116.97 95%HPD) in the two genes-based tree and 110.26 Mya (91.43-126.3 95%HPD) in the five genes-based tree. Again, these estimations were in agreement with previous studies Misof *et al.* (2014); Santos-Garcia *et al.* (2015).

The most studied whitefly species, the *B. tabaci* species complex, was part of the green cluster. Our estimations of the emergence time of the *Bemisia* genus and the *B. tabaci* species complex were similar to previous estimations by Santos-Garcia *et al*. (2015): 44.08 Mya (31.36-57.11 95%HPD) and 7.27 Mya (3.43-11.48 95%HPD) or 47.84 Mya (31.64-64.53 95%HPD) and 11.87 Mya (5.42-19.52 95%HPD), in the five genes- and two genes-based trees. Finally, the divergence between *B. tabaci* species MEAM1 and MED also agreed with previous estimates (Santos-Garcia *et al*., 2015). Taken together, although topological differences existed between the two trees, the convergence of their divergence time estimates supports their robustness.

### Tracking the origin of genomic instability in *Portiera*

*Portiera* from *B. tabaci* lacks the DNA polymerase III proofreading subunit (*dnaQ*). This absence seems ro be correlated with the massive rearrangements, large intergenic regions, and repetitive sequences (especially microsatellites) present in this *Portiera* lineage (Sloan and Moran, 2013; Santos-Garcia *et al*., 2015). To locate the evolutionary point in which *Portiera* lost *dnaQ* and genome stability begans, we screened our samples for the presence of *dnaQ* and four possible gene order configurations. The two configurations *groEL*-*rpsA* (A) and *leuC* -*leuD* (B) are ancient ones and are shared between *Aleurodicus* and *T. vaporariorum*. The other two rearrangements, *lepA*-*groEL* (A_*Bt*_) and*secA*-*leuC* (B_*Bt*_), were found so far only in the *B. tabaci* complex (Figure 3).

**Figure 3:**
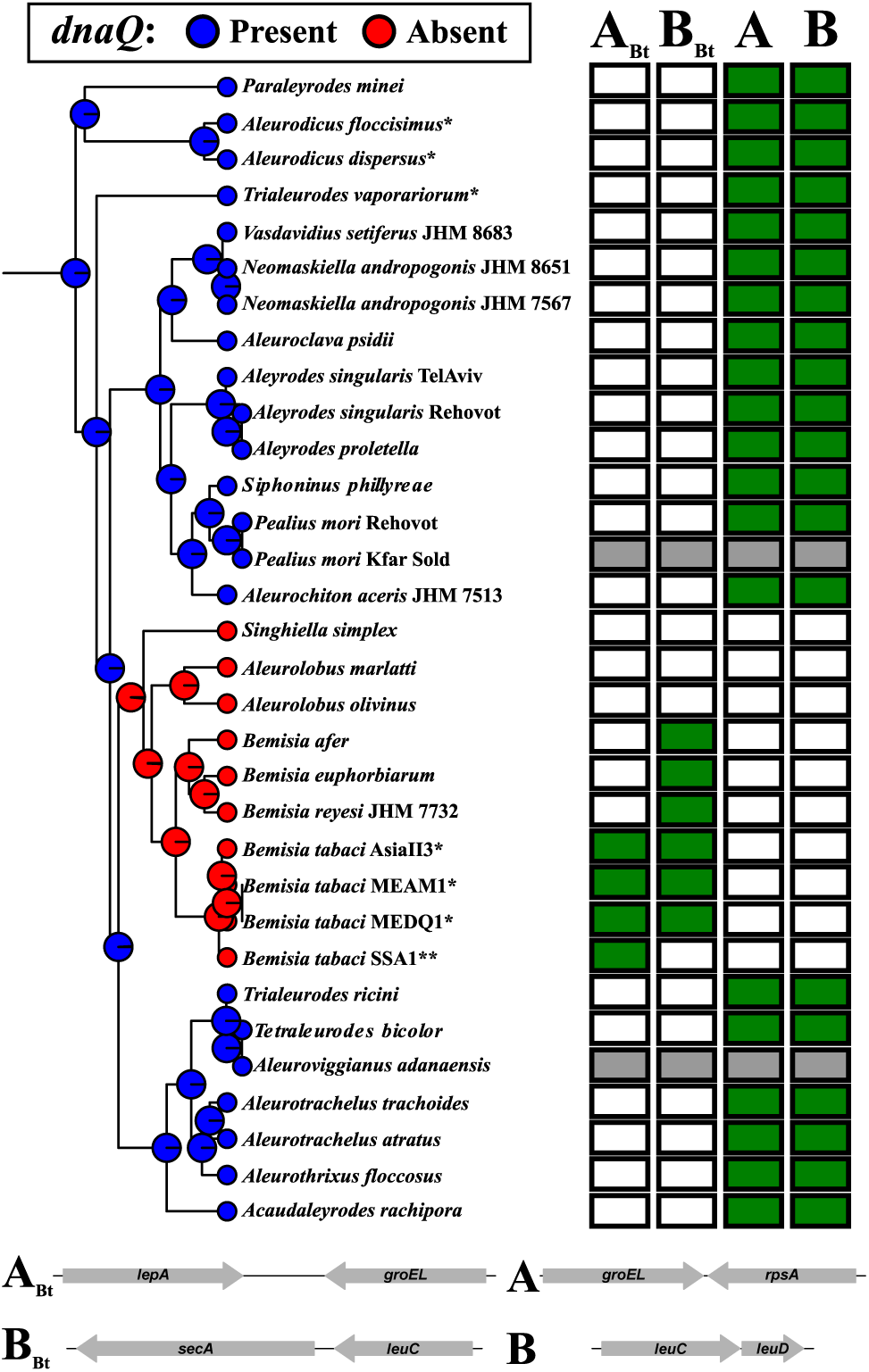
Summary of the screening for the *dnaQ* gene presence or absence and the gene rearrangements. Ancestral state inference was estimated using the five *Portiera* genes-based tree (left). Pie charts at the nodes represent the posterior probability for the presence (blue) or absence (red) of *dnaQ*. Note that all nodes have the probability of 1. The matrix represents the gene rearrangement amplification results (right). The letters above the matrix indicate the four possible rearrangements that were tested. Letters without index refer to the ancestral gene order found in *Portiera* of *Aleurodicus* and *T. vaporariorum*. Letters with the sub-index *Bt* refer to the gene order found in *Portiera* of *B. tabaci*. Green filled squares represent successful and validated amplifications while gray filled squares indicate that the gene order rearrangements were not tested but the ancestral one (A and B) is assumed. *: full genome available. **: no transcripts containing the B_*Bt*_ region were obtained.

We were able to amplify *dnaQ* of *Portiera* from all species tested except for *Aleurolobus olivinus, Aleurolobus marlatti, Singhiella simplex, Bemisia afer, B. euphorbiarum*, and *B. reyesi*. The first three genera species form a monophyletic clade together with *Bemisia*. Based on our ancestral state reconstruction using the five genes-based tree, it is highly likely (posterior probability of 1) that the most recent common ancestor (MRCA) of this clade also lacked a functional *dnaQ* gene (Fig 3). In addition, when using the two genes-based tree, it is probable that the MRCA of the *Singhiella*-*Bemisia* clade also lacked a functional *dnaQ* (0.62 posterior probability) (Figure S1). Due to too much uncertainty, we could not resolve the presence/absences of *dnaQ* in deeper nodes. Still, under a maximum parsimony scenario, *dnaQ* is likely to be present in the genome of *Portiera* from all whiteflies, with the exception of the *Singhiella*-*Bemisia* clade.

*Portiera* of all species outside the *Singhiella*-*Bemisia* clade also presented the ancestral gene order (rearrangements A and B) (Fig 3). *Portiera* of *Bemisia* species outside the *tabaci* species complex only presented the B_*Bt*_ rearrangement, suggesting a possible different order in region A. Although were unable to recover any transcript containing the B_*Bt*_ region in the *B. tabaci* SSA1 species, this species seems to be syntenic to other *B. tabaci* species (Figure S2). In addition, we could not amplify the ancestral or modified A and B regions in both of the *Aleurolobus* species and *Singhiella simplex*, raising the possibility that several gene rearrangements took place in the A and B regions during the evolution of *Portiera* in the *Singhiella*-*Bemisia* clade (Fig 3).

### The genomic and metabolomic characterization of *Portiera* from *Singhiella simplex*

To further elucidate the origin of *Portiera* genome instability and its putative effects on functionality, we sequenced the genome of *Portiera* from the most basal species in the *Bemisia* clade, the fig whitefly *Singhiella simplex*. As explained in length in the “*Materials and Methods*” section, the sample also contained individuals of the mulberry whitefly *Pealius mori*, which share some host-plants with *S. simplex*. As we were able to classify and recover complete *Portiera* and mitochondrial genomes from both *S. simplex* and *P. mori*, there was no effect to the accidental mixing on our further analyses. The genome of *Portiera* from *S. simplex* was recovered as 9 contigs (Table 1), all ending in repetitive sequences. It is the largest *Portiera* genome described so far, being 134Kb larger than that of *Portiera* of *B. tabaci* (Table 1). The number of coding genes was similar in *Portiera* of *S. simplex* and *B. tabaci*, which suggests that the genome expansion in *Portiera* of *S. simplex* is due to an increase in the size of the intergenic regions, which account for 40% of the genome. The genome of *Portiera* from *S. simplex* presented the lowest coding density (59.6%) and the highest number of direct (23) and inverted repeats (17) among all analyzed endosymbionts genomes (Table 1). As was already “suspected” from the PCR amplification results, the *dnaQ* gene was found to be pseudogenized in *Portiera* of *S. simplex*.

**Table 1:**
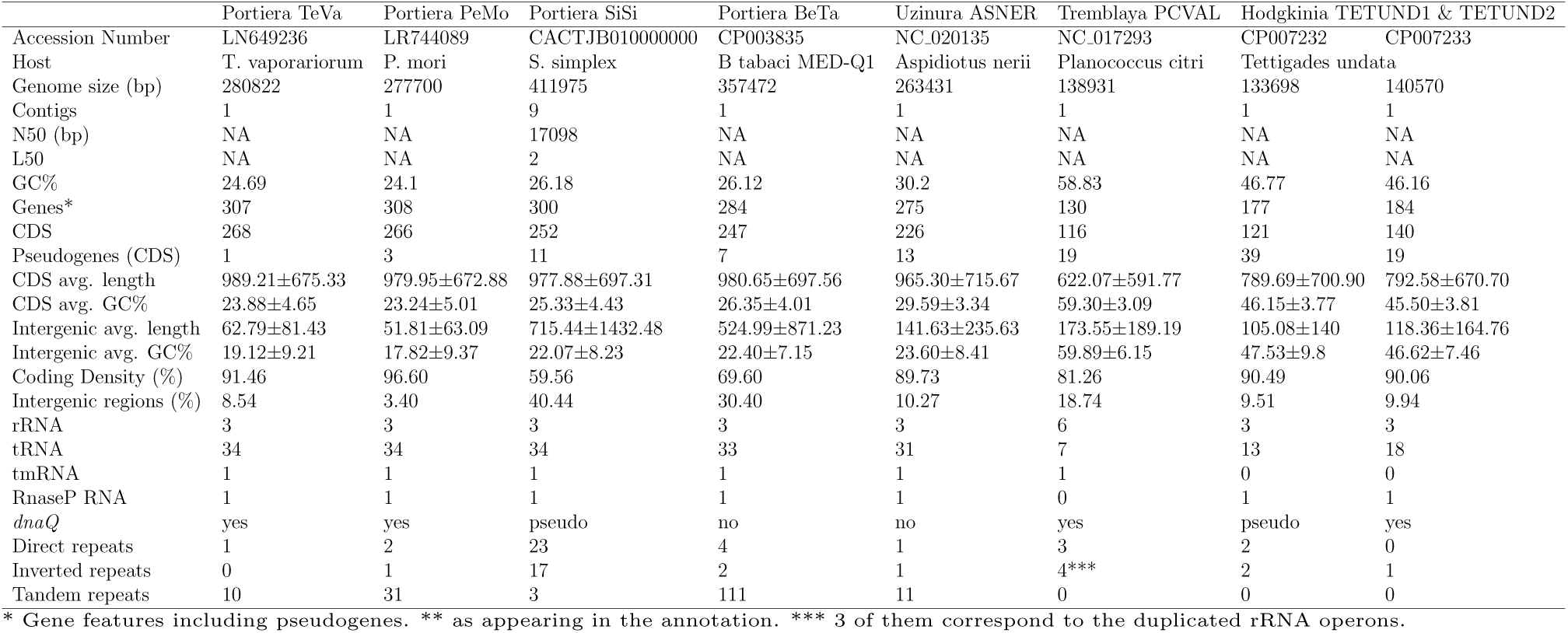
General genomic features of *Portiera* and other endosymbionts lacking *dnaQ*, presenting large intergenic regions or genome instability.

Synteny evaluation analysis, based on 235 Orthologous Clusters of Proteins (OCPs) (Figure S3), indicated that *Portiera* of *S. simplex* presents a different genomic architecture when compared both to the ancestral *Poritera* or to *Portiera* of *B. tabaci* gene order (Figure 4A). Still, textitPortiera of *S. simplex* presented a high degree of microsynteny (e.g. operons) with the other *Portiera* genomes. The *dnaQ* pseudogene is located in a region that has suffered different rearrangements and an expansion of the intergenic regions (Figure 4B). Comparisons to other *Portiera* genomes and different obligatory endosymbionts present in mealybugs, scale insects, and cicadas (Figure S4), indicated that endosymbionts lacking a functional *dnaQ* present extended intergenic regions (Kruskal-Wallis test, df = 8, p-value *<* 2.2^*e*−16^ and pairwise Wilcoxon test with Benjamini–Hochberg FDR, Figure S4).

**Figure 4:**
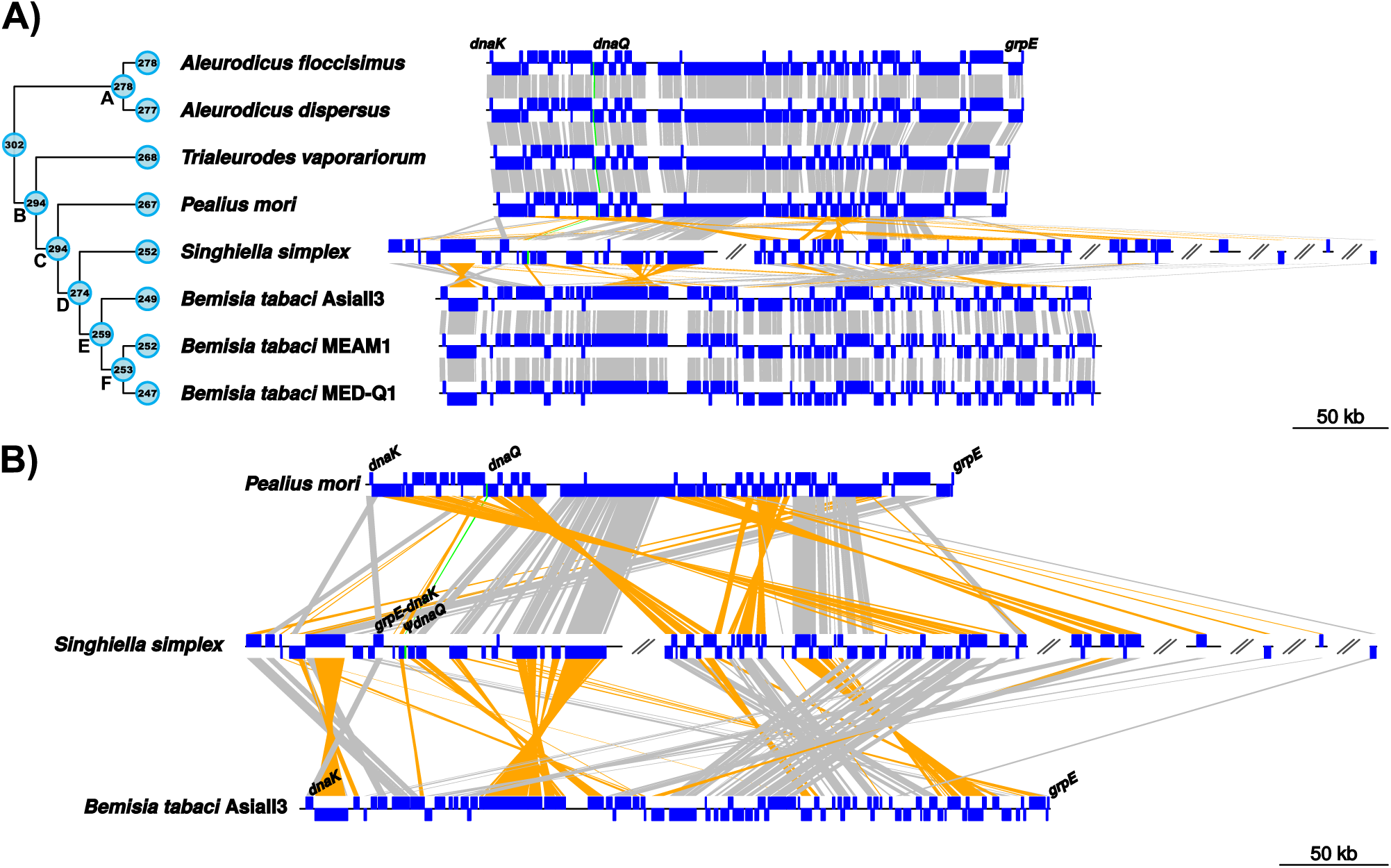
*Portiera* genomes syntenic comparisons based on 230 single copy core genes. **A)** Cladogram summarizing *Portiera* phylogenetic relationships based on the species tree obtained as part of the *OrthoFinder* pipeline. Filled circles at the nodes represent the number of coding genes estimated to be present in the most common recent ancestor (MRCA) using COUNT. Filled circles at the leaf tips represent the number of coding genes in each *Portiera* genome. Letters at the nodes list the MRCAs to allow comparisons with Table 2. *Portiera* genomes are represented linearly with blue boxes representing syntenic genes in the direct strand (upwards) or in the complementary strand (downwards), gray lines connect genes in the same strand, yellow lines connect genes in different strands, twisted lines indicate inversions. The green line highlights the position of functional and non-functional (*ψ*) *dnaQ* genes. For *Portiera* of *S. simplex*, only contigs containing core genes are represented (seven from nine contigs). **B)** Magnification of syntenyc comparisons between *Portiera* of *P. mori, S. simplex*, and *B. tabaci* AsiaII3.

Metabolically, the genome of *Portiera* of *S. simplex* is close to *Portiera* of *B. tabaci*. It has incomplete pathways for lysine but can still produce arginine. It has lost the ability to produce tryptophan (the *trpF* gene is absent) and also lacks the aminoacyl-tRNA synthetases *metG* and *alaS* (also lost *Portiera* of *T. vaporariorum, P. mori*, and *B. tabaci*) and *trpS* (lost in *Portiera* of *B. tabaci*) (Table 2). The tRNA^*Ile*^-lysidine synthetase *tilS*, responsible for avoiding miss-charging of methionine instead of isoleucine, was found to be uniquely pseudogenized in *Portiera* of *S*.*simplex*. In addition, a large gene-loss event (18 genes) occurred in MRCA of the *Singhiella*-*Bemisia* clade (Table 2 and Figure 4A). This event included the loss of *dnaQ* and other six genes related to the DNA replication and repair machinery (Table 2).

**Table 2:**
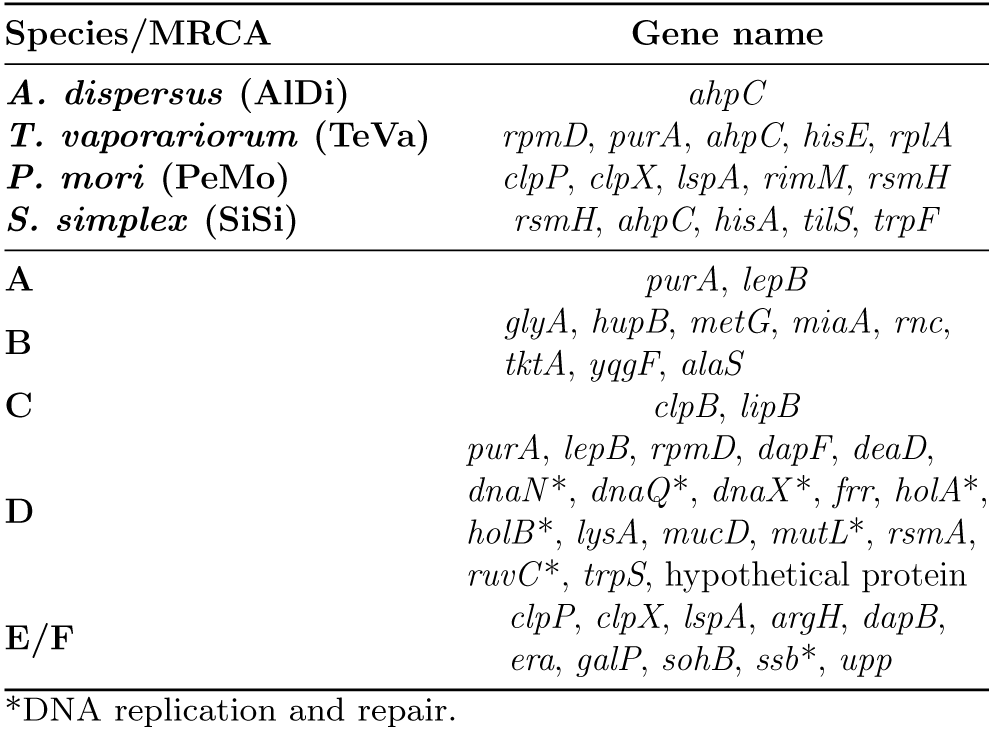
Gene losses in *Portiera* lineages and their MRCAs (nodes from Figure 4).

### Comparative Molecular Evolution of *Portiera* lineages

We estimated the ratio of synonymous (*S*) and non-synonymous (*N*) substitutions per site (*dS* and *dN*) and their omega ratio (*ω* = *dN/dS*) in 232 single-copy genes shared between the *Portiera* lineages of six whitefly species: *A. dispersus, A. floccissimus, T. vaporariorum, P. mori, S. simplex* and *B. tabaci* (MEAM1). After filtering, 158 orthologous shared genes were kept. To obtain the *S* and *N* per site per year (*dS/t* and *dN/t*), the values were divided by the lineage divergence time: 19.64 Myr (Million years) for *Aleurodicus*, 111.29 Myr for *Trialeurodes*, 99.98 Myr for *Pealius*, and 71.34 for *Singhiella*-*Bemisia* (Figure 1). *dS/t* and *dN/t* are normalized values and allow comparisons between lineages (the branch leading to a specific Portiera genome). For analysis, we used the chronogram that was based on five *Portiera* genes as it presented higher internal nodes support.

Our analyses indicated that the *Portiera* lineages evolve at different *dS/t* (KruskalWallis test, p-value *<* 2.2e-16) (Figure 5A). *Portiera* of *B. tabaci* was the fastest-evolving lineage, followed by *Portiera* of *S. simplex. Portiera* of *P. mori* was the slowest evolving lineage (Table 3). Also, *dN/t* values showed statistical differences among *Portiera* lineages (Kruskal-Wallis test, p-value *<* 2.2e-16) (Figure 5B). Similar to *dS* rates, *Portiera* of *B. tabaci* and *S. simplex* were the fastest evolving lineages while *Portiera* of *P. mori* was the slowest evolving lineage (Table 3).

**Table 3:**
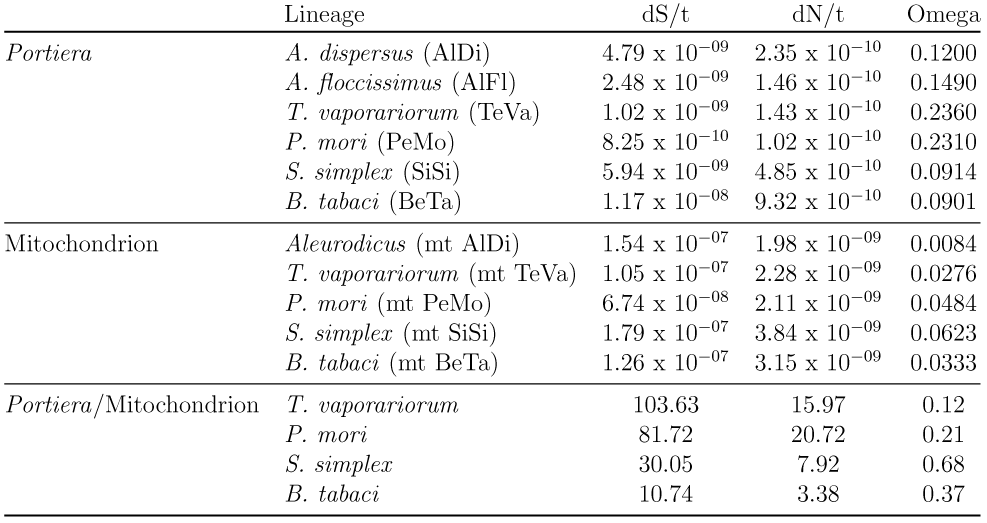
Average nucleotide substitutions per site per year and *ω* ratios for *Portiera* and mitochondrial lineages.

**Figure 5:**
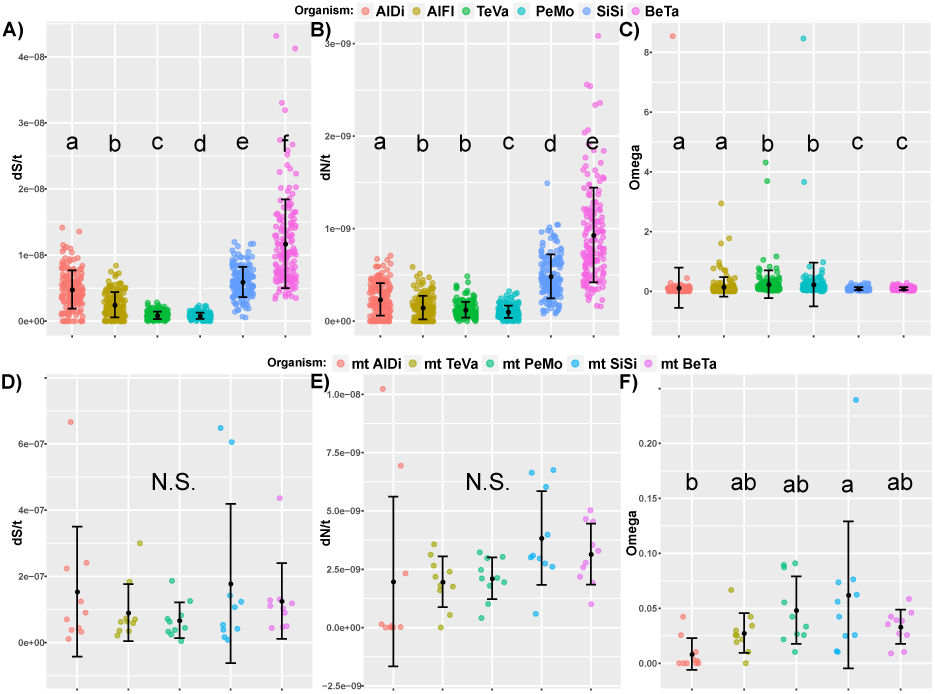
Synonymous (**A**) and nonsynonymous (**B**) substitutions per site per year and their *ω* ratios (**C**) estimated for 158 core shared genes between *Portiera* lineages of six whitefly species. Different letters indicate significant statistical differences between lineages (non-parametric Kruskal-Wallis and Wilcoxon post-hoc pairwise tests). Synonymous (**D**) and non-synonymous (**E**) substitutions per site per year and *ω* ratios (**F**) estimated for 10 full mitochondrial genes from six whiteflies species. N.S. = No significant difference. Different letters indicate significant statistical differences between lineages (one-way ANOVA and Tukey’s post-hoc test).

Next, we tested if the six Portiera lineages differ in the selection forces that act on their genomes by comparing their *ω* ratios (Figure 5C). Three statistically significant groups were observed (Kruskal-Wallis test, p-value *<* 2.2e-16): *Portiera* of *B. tabaci* and *S. simplex* had the lowest *ω* values, *Portiera* of *A. dispersus* and *A. floccissimus* presented intermediate *ω* values, and *Portiera* of *T. vaporariorum* and *P. mori* had the highest *ω* values. Most *ω* values were close to zero indicating a strong effect of purifying selection in almost all tested genes. Still, we detected 18 genes presenting signatures of relaxed/adaptive selection (*dS >* 0, *ω* ≤ 1 & *ω* ≥ 10) in *Portiera* of *T. vaporariorum* (9 genes), *A. floccissimus* (4), *P. mori* (3), and *A. dispersus* (1) (Table S4). Some of these genes were found to be related to amino acid biosynthesis (*hisH, leuC, trpC, gatC*), aminoacyl-tRNA synthetases (*sufS* and *cysS*), or to energy metabolism (*atpB* and *cyoD*).

Because the absence of host genomic/transcriptomic resources for a sufficient number of whitefly species, we estimated the mitochondrial nucleotide substitution rates as a proximation of nucleotide substitution rates in the host genomes (Figure 5D-F). Since the mitogenome of *A. floccissimus* is still not available, we calculated the *dS/t* and *dN/t* values of the *Aleurodicus* lineage using only the mitogenome of *A. dispersus* (dividing the values by 129.35 Mya, the estimated time when the split between the Aleurodicinae and the Aleyrodinae families occurred). Only 12 genes were included in the analysis since mitogenomes annotation was not consistent and *COXII* was filtered out. The mitochondrial lineages presented similar *dS/t* and *dN/t* values (One-way ANOVA, p-value *>* 0.2) (Figure 5D and E and Table 3). The *ω* values differed among lineages, being *S. simplex* and *A. dispersus* mitogenomes the ones with the higher and lowest values, respectively (One-way ANOVA, p-value *<* 0.02 and Tukey’s post-hoc test) (Figure 5F). Yet, in all lineages nearly all *ω* values were below 0.1, indicating a strong effect of purifying selection. Attending to the *dS/t* ratio between mitochondria and *Portiera*, the former is evolving faster. These ratios vary from ten in *B. tabaci* to 100 in *T. vaporariorum*.

## 4 Discussion

### *Portiera* as a valuable phylogenetic resource for studying whitefly taxonomy

Unlike taxonomical research in other insect groups that rely on adult morphology for classification, the current taxonomy of white-flies is mostly based on the morphology of the nymphal stages. These stages present plasticity in many morphological traits that respond to various abiotic and biotic environments including the identity of the plant host (Charles, 2010; Manzari and Quicke, 2006), eliminating in many cases the possibility of identifying a clear objective criterion. This has led to relatively high number of inaccuracies and mis-assignments in the group taxonomy (Manzari and Quicke, 2006). For example, the *Dialeurodes* genus, which is one of the most prolific and studied groups, lacks clear taxonomical keys. As a result, the genus is currently divided into at least five major groups, each containing several genera (Jensen, 1999, 2001).Additional support to the current problematic status of whitefly taxonomy comes from two complementing large-scale studies. In the first, an extensive cladistic analysis suggested that around half of the 117 Aleyrodinae genera analyzed were not monophyletic (excluding monobasic genera) (Manzari and Quicke, 2006). The second study used puparial morphological characters from all 20 Aleurodicinae genera and DNA sequences from nine Aleurodicinae genera, but recovered only 60% and 14% of the genera as monophyletic, respectively (Charles, 2010). Taking all above in consideration, it is quite safe to state that whitefly taxonomy can significantly benefit from the development of complementary classification frameworks, especially those using molecular data.

The current most popular molecular framework uses *mtCOI* sequences for species “barcoding”. In whiteflies, two *mtCOI* - dependent barcoding approaches are used. The first approach uses the 5’ region of the *mtCOI* gene. This region was used by the Consortium for the Barcode of Life (BOLD) initiative (www.barcodeoflife.org) which currently includes 1062 putative species (Barcode Index Number System clusters) and 102 identified species. The second approach, which is traditionally used in *Bemisia tabaci* research, targets the 3’ region of the *mtCOI* gene (Frohlich *et al*., 1999; De Barro *et al*., 2011). Using a modified 5’ *mtCOI* barcoding approach (cocktail of primer sets based on the LCO1490 and HCO2198 primers), the phylogenenetic tree published by Dickey *et al*. (2015) failed to recover the two whitefly subfamily lineages and only few nodes were well supported. Interestingly, the *Singhiella*-*Aleurolobus*-*Bemisia* clade was recovered with reasonable support. Similarly, Ovalle *et al*. (2014) used the 5’ *mtCOI* to explore the relationship between 8 genera (20 taxa). Again, while deeper nodes were not fully resolved, more recent nodes were well supported. This raises the possibility that the use of *mtCOI* gene sequences might be a powerful framework for whitefly barcoding (species identification) but a less accurate approach for establishing genus boundaries and genera relationships in the superfamily, as already was proposed for the Aphidoidea (Coeur d’Acier *et al*., 2014).

The low success of using mitochondria gene sequences for establishing inter-genus relationships in whiteflies, could be an effect of the mitogenome mutation rate, which can be more than 20-times higher that of nuclear genes (Allio *et al*., 2017). Moreover, whitefly mitogenomes in particular, present several translocations and evolve faster than other hemipteran insects (Thao *et al*., 2004; Song *et al*., 2012). Therefore, it is not surprising that we found that many mitochondrial genes show saturation at their synonymous sites, suggesting a compromised phylogenetic signal. For example, the widely used *mtCOI* gene shows saturation in many third codon positions (Table S3) (Song *et al*., 2012). Therefore, the third codon position should be excluded from phylogenetic analysis, reducing the effective length of the alignment. Although full mitogenomes can be used to resolve deeper relationships, the mitochondrial genes to be included should be selected carefully to avoid saturated ones. Even so, this approach seems to be unworkable in cases of large screening. A second problem we detected when using the *mtCOI* universal primers is their relatively low (around 58%) amplification success rate.

### The pros and cons of using *Portiera* gene sequences for studying whitefly taxonomy and phylogenetics

Several studies have proved the accuracy of using gene sequences of primary endosymbionts for reconstructing their host evolutionary history (Martinez-Torres *et al*., 2001; Jousselin *et al*., 2009; Nováková *et al*., 2013; Meseguer *et al*., 2015; Hall *et al*., 2016; Meseguer *et al*., 2017). Here, we found several additional advantages for the use of *Portiera* gene sequences in inferring the phylogenetic relationships among whiteflies. First, in contrast to *mtCOI* amplicons, all designed *Portiera* primers had an almost perfect amplification success except for the *rpoD* set that failed to amplify one sample. Second, the specific targeting of *Portiera* genes is by itself a diagnostic tool that allows to differentiate whiteflies from similar insects (e.g., psyllids nymphal stages) and to discriminate between the two whitefly subfamilies. Discrimination is possible because *Portiera* from the Aleurodicinae subfamily presents two specific insertions in the *23S* rRNA gene (Thao and Baumann, 2004b). Third, targeting *Portiera* genes is especially useful when studying parasitized samples (Table S2), as the use of universal *mtCOI* primers is in this case problematic. Fourth, *Portiera* is evolving slower than the mitogenome of whiteflies 3. Hence, *Portiera* genes usually do not show phylogenetic signal saturation. This makes the use of *Portiera* gene sequences more adequate for solving inter-genera relationships and deeper nodes than the *mtCOI* gene sequence. On the other hand, *Portiera* gene sequences seem to be limited in their ability to resolve recent speciation events or minor differentiations between cryptic species. It is also likely that at the intra-species level (populations), fast-evolving genes, as the *mtCOI*, will perform better. Last but not least, there are several *Portiera* genomes available, which allows targeting different sets of genes.

### The problematics and complexity of whitefly taxonomy

Some genera included in our analysis previously presented phylogenetic incongruence Manzari and Quicke (2006). Among them, *Aleurolobus, Aleuroplatus, Aleurotrachelus, Bemisia, Dialeurodes, Tetraleurodes*, and *Trialeurodes* had more than one representative in our analysis. Therefore, we tried solving previously observed phylogenetic conflicts by applying the *Portiera* phylogenetic framework established. In the case of *Aleuroplatus*, our results seems to support the paraphyly of these genera. Also, our phylogentic reconstruction supports the monophyly of the*Dialeurodes* and *Tetraleurodes* genera (Manzari and Quicke, 2006) if *Tetraleurodes bicolor* and the two unidentified *Dialeroudes* samples clustering with *Trialeurodes* are not taken into account. In addition, our analysis confirmed the relationship between *Aleuroclava* and *Dialeurodes*. Based on their early divergence, which fits more a separation time period between genera, both *Dialeurodes citri* and *Dialeurodes hongkongensis* are likely to belong to two different genera, with the latter probably assigned to the *Gigaleurodes* Jensen *(1999); Manzari and Quicke (2006)*.

*Aleurolobus, Aleurotrachelus*, and *Bemisia* were monophyletic in our study and also in previous analysis (Ovalle *et al*., 2014), in contrast to their paraphyletic status reported in Manzari and Quicke (2006). The only *Aleurothrixus* species in our dataset grouped with the *Aleurotrachelus* species, suggesting that this genus could be synonymized with the latter. Although never grouped together (Manzari and Quicke, 2006), our work clearly supports the relationship between the *Aleurolobus* and *Bemisia* genera, as was previously suggested (Ovalle *et al*., 2014). Interestingly, *Singhiella simplex* established a monophyletic clade together with *Aleurolobus* and *Bemisia* in our work. This finding disagrees with previous studies that suggested relationship between the *Singhiella* and *Dialeurodes* genera (Jensen, 2001; Manzari and Quicke, 2006). The possible existence of a *Singhiella*-*Aleurolobus*-*Bemisia* clade is also supported by *mtCOI* phylogenetic analysis (Ovalle *et al*., 2014; Dickey *et al*., 2015). However, while morphological classification of *Aleurolobus* species is straightforward (Manzari and Quicke, 2006), it could be possible that *Shingella simplex* has been erroneously assigned to the *Shingella* genus.

Several authors have tried to classify the Aleryrodinae into different tribes (reviewed in Manzari and Quicke (2006)). Despite our limited sampling, our results support some of those classifications. Based on their monophyly and the genome characteristics of their *Portiera* lineage (see below), we propose that the genera *Singhiella, Aleurolobus*, and *Bemisia* belong to the Aleurolobini tribe. The Aleurocanthini tribe was suggested to include the *Aleurocanthus, Aleurotrachelus*, and *Pentaleyrodes* genera. As support for this tribe, we recovered a monophyletic clade including the *Aleurotrachelus* and *Aleurothrixus*. We found that the Neomaskellini tribe includes both the *Neomaskellia* and *Vasdavidius* genera. Two from three *Tetraleurdes* species were recovered as monophyletic, suggesting that the Tetraleurodini tribe is likely to be correct. The tribes Aleyrodini and Diaeleurodini are the largest but also the most controversial. In these tribes, we can only confirm the Diaeleurodini membership of both *Dialeurodes* and *Aleuroclava*.

Finally, we would like to comment on two inconsistencies in our data. The almost identical *Portiera* sequences of *Aleurovig-gianus adanaensis, Tetraleurodes bicolor*, and *Trialeurodes ricini* could be due to cross-contamination or possible adult miss-classification (e.g. adult near puparias from different species). If *Trialeurodes ricini* is not considered, our results support the possibility that *Trialeurodes* is a monophyletic genus (Manzari and Quicke, 2006). Contrarily, the identical *Portiera* sequences of *Aleyrodes singularis* and *Aleyrodes prolotella* are likely to reflect a recent speciation event with some hots-plant overlap (Martin *et al*., 2000). *Aleyroides* cross-contamination is unlikely as only *Aleyrodes singularis* has been found in Israel and gDNA extractions and amplicons amplification were performed separately in time for each species.

### *Paraleyrodes*: a living fossil?

Described at first as *Aleyrodes perseae*, the nymphal stages of the genus *Paraleyrodes* present typical Aleurodicinae morphological characters, such as subdorsal compound pores or legs with apical claws (Quaintance, 1909). However, adults present morphological characters typical of Aleyrodinae, such as small body size and single-vein wings Quaintance (1909); Martin (1996, 2007). Interestingly, the *Paralyerodes* genus presents median ocellus, an ancestral character described in Cretaceous taxa (Drohojowska and Szwedo, 2015). Our analysis supports the inclusion of the *Paraleyrodes* genus inside the Aleurodicinae subfamily based on its ancient origin and the presence of the *23S* rRNA insertions common to the Aleurodicinae subfamily (Thao and Baumann, 2004b). Our estimates overlap with the calibration point used, suggesting that *Paraleyrodes* genus originated in the Lower Cretaceous (100.5-145 Mya). Therefore, *Paraleyrodes* can be considered a long-enduring extant taxon, which may explain the retention of the middle ocellum and the mixture of morphological characteristics of both Aleyrodidae subfamilies. Although speculative, it also could be possible that other hard-to-assign Aleurodicinae genera, such as *Aleuroctarthrus* (presents medium ocellus) or *Palaealeurodicus* (does not present clawed legs), are indeed long-enduring taxa (Martin, 2008). Using cladistic and molecular analysis, these two genera were closely related to *Paralyerodes* (Charles, 2010). *In addition, Palaealeurodicus* was placed as basal to all Aleurodicinae based on four mitochondrial genes (Charles, 2010). Therefore, *Paraleyrodes* can be considered as sister taxon of *Palaealeurodicus*, which diverged before the radiation of the *Aleurodicus* genus (Charles, 2010). Identifying such kind of long-enduring taxa could be an invaluable resource for understanding the evolution of the whitefly superfamily.

### Long-standing endosymbionts present almost static genomes

Adaptation to the intracellular lifestyle has a significant impact on bacterial symbionts. Metabolic redundancy between the host and the endosymbiont promotes the dependency of the later on the former intracellular environment (Morris *et al*., 2012). Moreover, vertical transmission drastically reduces the endosymbiont effective population size (*N*_*e*_) and the chances to acquire new genetic material, eventually leading to the generation of asexual populations. The combined effects of vertical transmission and intracellular lifestyle promote the accumulation of deleterious mutations that are otherwise pruned by selection in larger *N*_*e*_, which can lead to massive loss of genes (Moran, 1996; Toft and Andersson, 2010; Wernegreen, 2015). The outcome of the process, known as the Muller’s Ratchet (Moran, 1996), is an endosymbiont that harbors a highly reduced genome, with small intergenic regions and very few repetitive sequences (Toft and Andersson, 2010; Wernegreen, 2015). Common conserved elements include genes that are essential for producing the host requirements and a minimal informational and translational machinery required for cell maintenance (Moran and Bennett, 2014).

As a consequence of the reduced or absent replication and recombination machinery, and the minimal presence of repetitive sequences, long-standing endosymbionts present almost static genomes (Moran and Bennett, 2014). For example, only few inversions were detected in endosymbionts that have been co-diverging with their host for more than 100 Myr (Patiño-Navarrete *et al*., 2013; Chong *et al*., 2019). *Portiera* of whiteflies is not different from other long-standing endosymbionts and usually harbor “classical” reduced and static genomes (Sloan and Moran, 2013; Santos-Garcia *et al*., 2015). One exception to this “rule” is *Portiera* of *B. tabaci*, which presents a genome with large intergenic regions, extensive rearrangements, and abundance of repetitive sequences (Sloan and Moran, 2013; Moran and Bennett, 2014; Santos-Garcia *et al*., 2015).

### Genome instability in *Portiera* of the tribe Aleurilobini

The genome of *Portiera* from *B. tabaci* presents one of the most reduced sets of DNA replication and repair genes among known long-standing P-endosymbionts, including the loss of the *dnaQ* gene which encodes the DNA polymerase proofreading subunit (Moran and Bennett, 2014). This loss has been linked to the uncommon extensive genome re-arrangements, inversions, abundance of repeated sequences, large intergenic regions, and accelerated evolution found (Sloan and Moran, 2013; Santos-Garcia *et al*., 2015). Our findings here suggest that the massive loss of DNA replication and repair genes is not restricted to *B. tabaci*, but shared by all other members of the *Singhiella*-*Aleurolobus*-*Bemisia* clade (hereafter the tribe Aleurilobini for simplicity). Therefore, it is quite probable that *dnaQ* was pseudogenized in the last common ancestors of the tribe, more than 70 Mya.

So far, only three genomes displaying long intergenic regions, genome instability, or the lack of functional *dnaQ* have been sequenced from other long-standing endosymbionts: *Uzinura diaspidicola, Tremblaya princeps*, and *Hodgkinia cicadicola* (Moran and Bennett, 2014; Van Leuven *et al*., 2014; López-Madrigal *et al*., 2015; Łukasik *et al*., 2018). Only a single genome of *U. diaspidicola* is currently available, and therefore, it is not clear if this endosymbiont which lacks *dnaQ* also displays significant genome instability. Relative to the genomes of *Portiera* from *B. tabaci* and *S. simplex*, the genome of *U. diaspidicola* presents lower number of repeated sequences and smaller intergenic regions. One explanation to this could be the conservation of the *mutL* gene in *U. diaspidicola* (Moran and Bennett, 2014). The enzyme MutL, together with MutS, is part of the mismatch repair system that corrects mismatch events that are produced by base miss-incorporation and polymerase slippage (Rocha, 2003). The genome of *Tr. princeps* presents genome instability signatures such as long intergenic regions, gene conversions, and the presence of several direct/indirect sequence repeats (López-Madrigal *et al*., 2015). Still, the number of repeats and the length of the intergenic regions are smaller than in *Portiera* of *B. tabaci* or *S. simplex*. The inactivation of the recombination machinery in *Tr. princeps* has been proposed as a strategy to reduce the number of homologous recombination events and their deleterious consequences in highly reduced genomes (López-Madrigal *et al*., 2015). However, *Tr. princeps* has access to a complementing recombination machinery as it harbors the endosymbiont *Moranella endobia* which has an active recombination machinery (López-Madrigal *et al*., 2015). The presence of a functional *dnaQ* subunit in *Tr. princeps* and the possible access to a complementing recombination machinery suggest that the possible causes of genome instability in *Tr. princeps* are different from that of *Portiera*.

One of the most extreme cases of genome instability was reported in *H. cicadicola*. In some cicada genera, which usually have a long lifespan, *H. cicadicola* has been split into several lineages with different genomic content within the same insect. This enforces functional complementation between the lineages for normal growth (Van Leuven *et al*., 2014; Łukasik *et al*., 2018). Although the genomic architecture of *H. cicadicola* resembles that of *Portiera* presenting unstable genomes, there are major differences in their relationship with their hosts. While *Portiera* is essential for whiteflies, *H. cicadicola* is a co-primary endosymbiont in cicadas and has been replaced several times (Łukasik *et al*., 2018; Matsuura *et al*., 2018). Therefore, the selection forces acting on both endosymbionts could be very different: strong purifying selection in the case of *Portiera* while more relaxed, or even non-adaptive selection in the case of *H. cicadicola* (Łukasik *et al*., 2018).

Large intergenic regions can allow *Portiera* to better tolerate rearrangements while the expansion of repeated sequences can increase the chance of deleterious homologous recombination events (Sloan and Moran, 2013). Considering that the *Portiera* genome presents signs of gene conversion and recombination, it could be possible that long intergenic regions and intergenic repeats are selected in *Portiera* presenting unstable genomes to increase their resilience against deleterious mutations. Repeated sequences mostly accumulate at the intergenic regions and pseudogenes of *Portiera* from *B. tabaci* and *S. simplex* suggesting strong purifying selection at the gene level. However, it could be possible that recombination is also counter-selected in *Portiera* with unstable genomes. This could be the reason why *Portiera* lineages within the *B. tabaci* species complex are syntenic after, at least, 7 My of divergence and present low number of direct/indirect repeats compared to *S. simplex*. In the later, recombination seems still active. Therefore, it could be possible that after a period of genome instability and intergenic regions expansion, direct and indirect repeats are counter selected to favor more stable genomes.

In contrast, the location of tandem repeats in *Portiera* of *T. vaporariorum* (8 over 10) and *P. mori* (24 from 31) partially or completely overlap with coding genes. As sequence repeats in coding genes can cause gene inactivation and/or re-arrangements, their existence within genes of stable *Portiera* genomes can reflect the presence of a minimal, but functional, DNA repair machinery that allows a more relaxed purifying selection process. In fact, *Portiera* of *B. tabaci* and *S. simplex* showed the lowest *ω* values, indicating stronger purifying selection forces acting on their genomes. Taking together, it could be possible that increased resilience combined with a strong purifying selection force at the gene level, have helped to maintain *Portiera* away from extinction in the Aleurilobini tribe (Bennett and Moran, 2015).

### Possible processes affecting substitution rates in *Portiera*

There are clear differences in the substitution rates among *Portiera* lineages presenting a relatively similar set of recombination and repair genes. *Portiera* of *B. tabaci* substitution rates are close to those of two known accelerated endosymbionts: *Blochmania* (P-endosymbiont of ants) and *Buchnera* (P-endosymbiont of aphids) (Silva and Santos-Garcia, 2015). The substitution rates of *Portiera* from *S. simplex* fall in the range of free-living bacteria, as previously reported for *Portiera* of *Aleurodicus* and *Trialeurodes* (Santos-Garcia *et al*., 2015). *Portiera* of *P. mori* has substitution rates smaller than free-living bacteria and closer to those estimated for *Sulcia* (P-endosymbiont of cicadas), one of the oldest and slow-evolving endosymbiotic lineages described so far (Silva and Santos-Garcia, 2015).

One possible factor that can have an effect on the mutation rates of endosymbionts is their host generation time (Santos-Garcia *et al*., 2015; Silva and Santos-Garcia, 2015). There seems to be a positive correlation between the number of generations produced by the insect host and the *Portiera* substitution rates. For example, while *A. dispersus, A. floccissimus, T. vaporariorum*, and *B. tabaci* establish multiple generations nearly all year round, *S. simplex* and *P. mori* produce only around four *∼*generations per year (Table S2).

A second factor is related to the endosymbiont generation time (Santos-Garcia *et al*., 2015; Silva and Santos-Garcia, 2015). It has been reported that increased mutation rates and genome reduction are negatively correlated with cell growth Nishimura *et al*. (2017). This means that differences in gene losses can result in reduced metabolism in some lineages leading to slower growth rates. In the case of *Portiera* from *B. tabaci* and *S. simplex*, their gene content is more reduced than the rest of the *Portiera* lineages, despite having larger genomes. Moreover, in these two *Portiera* lineages, several gene losses (*dnaX, dnaN*, and *holAB*) are likely to affect the DNA polymerase III holoenzyme processivity. Longer replication time could produce slow-growing *Portiera* phenotypes, hence, longer generation times. Additionally, *Portiera* from *B. tabaci* has lost *ssb*, which plays a central role in DNA replication/repair (Shereda *et al*., 2008), increasing both the mutation rates and generation time compared to *Portiera* of *S. simplex*. It seems, therefore, that the increase in generation time can result both in reduced and increased mutational rates, depending on the mechanism involved. Reduced metabolic performance and longer replication time will decrease the mutational bias by restricting the number of *Portiera* cells divisions (Santos-Garcia *et al*., 2015; Silva and Santos-Garcia, 2015). On the other hand, increasing the replication time in the cell cycle due to reduced processivity of DNA polymerase III or reduced/lack of functionality of other important component in the replication machinery will promote higher mutation rates in spite of the prolonged generation time Nishimura *et al.* (2017).

Another factor that could affect the substitution rate of *Portiera* genomes is the possible complementation of the reduced DNA replication and repair machinery by host nuclear-encoded proteins. The host may exert this by two non-exclusive mechanisms. The first is using bacterial genes that have been horizontally transferred to the host nuclei and are targeted back to the *Portiera* cell. The second involvess the re-utilization of host mitochondria-targeted proteins (Santos-Garcia *et al*., 2014, 2015; Silva and Santos-Garcia, 2015). Although highly speculative, it is possible for the host cell to re-target the mitochondrial pol*γ*, which presents proof-reading activity, towards *Portiera*. Since *mutS* is present in all *Portiera* genomes, a combination of imported proteins and MutL-independent mismatch repair mechanism (Marti *et al*., 2002) can help to reduce the mutation rate in *Portiera* lineages, especially those presenting a reduced recombination and repair machinery. The possible involvement of host complementation mechanism is also supported by the finding that some host DNA replication and repair genes and several aminoacyl tRNA synthetases are over-expressed in *B. tabaci* ‘s bacteriocyte compared to the rest of the body (Mao *et al*., 2018).

### The symbiont or the egg: genome instability and bacteriocyte inheritance

Since the beginning of the research on insect symbiosis, it was clear that the whitefly superfamily presents a special mode of symbiont transmissio as whole maternal bacteriocytes are inherited by the offspring (Buchner, 1965). In *T. vaporariorum, Aleyrodes proletella, Aleurodes aceris*, and *Aleurochiton aceris* several bacteriocytes migrate to the oocyte (from five to ten depending on the species), entering through the future pedicel (Tremblay, 1959; Buchner, 1965; Szklarzewicz and Moskal, 2001). In contrast, in *B. tabaci, Bemisia aff. gigante*, and *Aleurolobus olivinus*, a single bacteriocyte enters the oocyte (Tremblay, 1959; Buchner, 1965; Coombs *et al*., 2007).

We are aware that the phylogenetic relationships of *B. aff. gigantea* are not completely resolved, but it does seem to be a sister clade of *Aleurolobus* and *B. afer* and distantly related to *B. tabaci* (Manzari and Quick*e, 2006)*. Therefore, according to the current published data, all the whitefly species that present a single-bacteriocyte mode of inheritance belong to only one phylogenetic group, the tribe Aleurilobini. The most parsimonious explanation suggests that the single-bacteriocyte mode of inheritance has evolved in the common ancestor of *Aleurolobus*-*Bemisia*, otherwise we will have to assume that it evolved multiple times in different species: *Aleurolobus olivinus, B. aff. gigantea* and *B. tabaci* (Tremblay, 1959; Coombs *et al*., 2007; Luan *et al*., 2016, 2018; Xu *et al*., 2019). Although we lack information on the mode of transmission of the bacteriocyte in *S. simplex*, there is a possibility that this species also presents the single-bacteriocyte mode of inheritance. If found to be true, it would suggest that the whole Aleurilobini tribe is likely to present this derived type of bacteriocyte inheritance.

There is an apparent relationship between the emergence of a single-bacteriocyte inheritance mode and the presence of *Portiera* lineages that show genomic instability. The evolution of a single-bacteriocyte inheritance mode could have had a big impact on *Portiera* evolution since it decreases the effective population size (*N*_*e*_) drastically compared to multi-bacteriocyte inheritance. The extremely low *N*_*e*_ probably intensified the effect of random genetic drift and accelerated the accumulation of deleterious mutations in *Portiera*. In addition, all the *Portiera* cells that are harbored in the same bacteriocyte are expected to present a homogenized allelic composition since recombination events, if happen, are limited to the cells inhabiting the same bacteriocyte. This implies a low chance of recovery of deleterious allele forms once they are established.

At the same time, the single-bacteriocyte inheritance mode also exerts strong purifying selection at both the bacteriocyte and *Portiera* levels each generation as offspring harboring a bacteriocyte or *Portiera* with deleterious mutations will probably sufferfrom severe fitness costs (Luan *et al*., 2018)). This is somewhat supported by the evidence that extant *Portiera* of the tribe Aleurilobini present moreover a stable gene content, with the massive gene loss events occurring only in their common ancestor. For instance, after approximately 70 Myr of divergence, only five and ten genes were lost from the *Portiera* genomes of *S. simplex* and *B. tabaci*, respectively.

Further research on the tribe Aleurilobini is required in order to determine what occurred first: the transition from the multi- to the single-bacteriocyte inheritance mode or the switch from stable to unstable genomic architecture of *Portiera*. In the first case, the evolution of a different mode of transmission could have triggered the DNA replication and repair machinery loss as purifying selection was not able to maintain them under very low *N*_*e*_ (Lynch, *2010)*. In addition, these losses may have been complemented by an overtake of some of their activities by the genome of the host cell (Santos-Garcia *et al*., 2014, 2015; Silva and Santos-Garcia, 2015; Mao *et al*., 2018). In the latter case, we should assume that *Portiera* lost their recombination and repair machinery as a consequence of a continuous genome degradation process (Bennett and Moran, 2015). This increased the chances for transmitting *Portiera* with deleterious mutations. A multiple-bacteriocytes inheritance mode results in the transmission of mixtures that can mask the presence of bacteriocytes harboring *Portiera* with deleterious mutations/variations. Instead, if single bacteriocytes are inherited, the *Portiera* presenting deleterious mutations will reduce the fitness of the new-born carrying them and they will be counter-selected. Therefore, the evolution of the single-bacteriocyte inheritance mode could have been a compensatory adaptation mechanism of the insect host to exercise an iron grip over *Portiera* transmission for ensuring the viability of its offspring (Campbell *et al*., 2018).

## 5 Conclusion

We argue here that the higher classification of whiteflies, at the levels of genera or tribe, requires reassessment. This problem is especially evident in the Aleyrodinae subfamily, which is highly diverse. The current use of nymph morphological characters is not sufficient, and molecular tools must be developed to support the classification of many species/genera in the superfamily. Our work brings evidence that the sequences of target genes of the primary endosymbiont of whiteflies, *Candidatus* Portiera aleyrodidarum, are a promising phylogenetic resource. They can be used to establish inter-genera relationships, can serve as diagnostic tools by themselves, and can help in the classification of problematic samples (even parasitized ones).

The ability of the established phylogenetic framework to solve standing inconsistencies in whitefly taxonomy also helped us localize a critical event in *Portiera* evolution, the appearance of genomic instability, which is very uncommon in primary endosymbionts. Our analyses suggest that the *Portiera* ancestor of the Aleurolibini tribe suffered a massive DNA replication and repair genes loss. This event was the trigger of the genomic instability shown by all extant *Portiera* lineages in the tribe. We hypothesize that the appearance of genomic instability is also related to the evolutionary switch made between multi- and single-bacteriocyte mode of inheritance.

## 6 Data deposition

Supplementary Material and Methods, Figures and Tables can be found at XXXX. Relevant scripts and files generated in this work will be available at https://figshare.com/s/8c7c9b4b1db43749ab6e. Sanger sequenced *Portiera* and *mtCOI* genes, Novaseq raw reads, and *Portiera* and mitochondrial assembled genomes have been deposited at the European Nucleotide Archive (ENA) under the project number PRJEB31657 and will be released after acceptance for publication.

## 7 Acknowledgments

The authors of this work want to honor the memory of Porf. Dan Gerling (1936–2016). This research was supported by the Israel Science Foundation grant Nos. 1039/12 to SM and 484/17 to EZ-F. DS-G was a recipient of the Golda Meir Postdoctoral Fellowship from the Hebrew University of Jerusalem. We wish to acknowledge Dra. Estrella Hernández Suárez and Dr. Hélène Delate for supplying whitefly samples.

## 8 Author contribution

DS-G and SM conceived the study. DS-G performed bioinformatics analysis and collected whitefly samples. NM-R performed molecular work and collected whitefly samples. DO supplied NHM whitefly samples and helped in whiteflies taxonomy. DS-G drafted the manuscript with inputs from SM. EZ-F and DO reviewed and corrected advanced versions of the manuscript.

## References

Allio R, Donega S, Galtier N, and Nabholz B. 2017. Large variation in the ratio of mitochondrial to nuclear mutation rate across animals: Implications for genetic diversity and the use of mitochondrial DNA as a molecular marker. Mol. Biol. Evol., 34(11):2762–2772.

Bankevich A, et al. 2012. SPAdes: a new genome assembly algorithm and its applications to single-cell sequencing. J. Comput. Biol., 19(5):455–77.

Bennett GM and Moran NA. 2015. Heritable symbiosis: The advantages and perils of an evolutionary rabbit hole. Proc. Natl. Acad. Sci., 112(33):10169–10176.

Boetzer M and Pirovano W. 2012. Toward almost closed genomes with GapFiller. Genome Biol., 13(6):R56.

Boetzer M, Henkel CV, Jansen HJ, Butler D, and Pirovano W. 2011. Scaffolding pre-assembled contigs using SSPACE. Bioinformatics, 27(4):578–579.

Bonfield JK and Whitwham A. 2010. Gap5-editing the billion fragment sequence assembly. Bioinformatics, 26(14):1699–1703.

Bouckaert R, et al. 2014. BEAST 2: A software platform for Bayesian evolutionary analysis. PLoS Comput Biol, 10(4):e1003537.

Boykin LM, Shatters RG, Rosell RC, McKenzie CL, Bagnall RA, De Barro P, and Frohlich DR. 2007. Global relationships of *Bemisia tabaci* (Hemiptera: Aleyrodidae) revealed using Bayesian analysis of mitochondrial *COI* DNA sequences. Molecular Phylogenetics and Evolution, 44(3):1306–1319.

Brown JK, Frohlich DR, and Rosell RC. 1995. The sweetpotato or silverleaf whiteflies: Biotypes of *Bemisia tabaci* or a species complex? Annual Review of Entomology, 40(1):511–534.

Buchner P. 1965. Endosymbiosis of animals with plant microorganisms, volume 7. John Wiley & Sons, Inc.: New York, New York. ISBN 978-0470115176.

Campbell BC, Steffen-Campbell JD, and Gill RJ. 1994. Evolutionary origin of whiteflies (Hemiptera: Sternorrhyncha: Aleyrodidae) inferred from *18S* rDNA sequences. Insect Mol. Biol., 3(2):73–88.

Campbell MA, et al. 2018. Changes in endosymbiont complexity drive host-level compensatory adaptations in cicadas. MBio, 9(6):8–11.

Castresana J. 2000. Selection of conserved blocks from multiple alignments for their use in phylogenetic analysis. Mol Biol Evol, 17(4):540–552.

Charles E. 2010. Systematics of whiteflies (Aleyrodidae: Aleurodicinae): Their distribution, phylogeny and relationship with parasitoids. Ph.D. thesis, Imperial College London.

Chen S, Zhou Y, Chen Y, and Gu J. 2018. fastp: an ultra-fast all-in-one FASTQ pre-processor. Bioinformatics, 34(17):i884–i890.

Chevreux, B, Wetter, T and Suhai S. 1999. Genome sequence assembly using trace signals and additional sequence information. Comput. Sci. Biol. Proc. Ger. Conf. Bioinforma., 99:45–56.

Chong RA, Park H, and Moran NA. 2019. Genome evolution of the obligate endosymbiont *Buchnera aphidicola*. Mol. Biol. Evol., 36(7):1481–1489.

Coeur d’Acier A, et al. 2014. DNA barcoding and the associated PhylAphidB@se website for the identification of European aphids (Insecta: Hemiptera: Aphididae). PLoS One, 9(6):e97620.

Conway JR, Lex A, and Gehlenborg N. 2017. UpSetR: an R package for the visualization of intersecting sets and their properties. Bioinformatics, 33(18):2938–2940.

Coombs MT, Costa HS, De Barro P, and Rosell RC. 2007. Pre-imaginal egg maturation and bacteriocyte inclusion in *Bemisia aff. gigantea* (Hemiptera: Aleyrodidae). Ann. Entomol. Soc. Am., 100(5):736–744.

Cryan JR and Urban JM. 2012. Higher-level phylogeny of the insect order Hemiptera: is Auchenorrhyncha really paraphyletic? Systematic Entomology, 37(1):7–21.

Csuos M. 2010. Count: evolutionary analysis of phylogenetic profiles with parsimony and likelihood. Bioinformatics, 26(15):1910–1912.

De Barro PJ, Liu SS, Boykin LM, and Dinsdale AB. 2011. *Bemisia tabaci* : a statement of species status. Annual review of entomology, 56:1–19.

De Vienne DM, Refrégier G, López-Villavicencio M, Tellier A, Hood ME, and Giraud T. 2013. Cospeciation vs host-shift speciation: Methods for testing, evidence from natural associations and relation to coevolution. New Phytol, 198(2):347–385.

Dickey AM, Stocks IC, Smith T, Osborne L, and McKenzie CL. 2015. DNA barcode development for three recent exotic whitefly (Hemiptera: Aleyrodidae) invaders in Florida. Florida Entomol., 98(2):473–478.

Drohojowska J and Szwedo J. 2015. Early Cretaceous Aleyrodidae (Hemiptera: Sternorrhyncha) from the Lebanese amber. Cretac. Res., 52:368–389.

Emms DM and Kelly S. 2019. OrthoFinder: phylogenetic orthology inference for comparative genomics. Genome Biol, 20(1):238.

Evans GA. 2007. The whiteflies (Hemiptera: Aleyrodidae) of the world and their host plants and natural enemies. Technical report, USDA/Animal Plant Health Inspection Service (APHIS).

Folmer O, Black M, Hoeh W, Lutz R, and Vrijenhoek R. 1994. DNA primers for amplification of mitochondrial cytochrome c oxidase subunit I from diverse metazoan invertebrates. Mol Mar Biol Biotechnol, 3(5):294–299.

Frohlich, Torres-Jerez, Bedford, Markham, and Brown. 1999. A phylogeographical analysis of the *Bemisia tabaci* species complex based on mitochondrial DNA markers. Molecular ecology, 8(10):1683–91.

Grimaldi D and Engel M. 2005. Evolution of the insects. Cambridge University Press, New York. ISBN 110726877X.

Guy L, Kultima JR, and Andersson SGE. 2010. genoPlotR: comparative gene and genome visualization in R. Bioinformatics, 26(18):2334–5.

Hall AAG, et al. 2016. Codivergence of the primary bacterial endosymbiont of psyllids versus host switches and replacement of their secondary bacterial endosymbionts. Environmental Microbiology, 18(8):2591–2603.

Hansen AK and Moran Na. 2014. The impact of microbial symbionts on host plant utilization by herbivorous insects. Molecular Ecology, 23(6):1473–1496.

Hsieh CH, Ko CC, Chung CH, and Wang HY. 2014. Multilocus approach to clarify species status and the divergence history of the *Bemisia tabaci* (Hemiptera: Aleyrodidae) species complex. Molecular Phylogenetics and Evolution, 76(1):172–180.

Jensen AS. 1999. Cladistics of a sampling of the world’s diversity of whiteflies of the genus *Dialeurodes* (Hemiptera : Aleyrodidae). Ann. Entomol. Soc. Am., 92(3):359–369.

Jensen AS. 2001. A cladistic analysis of *Dialeurodes, Massilieurodes* and *Singhiella*, with notes and keys to the Nearctic species and descriptions of four new *Massilieurodes* species (Hemiptera: Aleyrodidae). Syst. Entomol., 26(3):279–310.

Jousselin E, Desdevises Y, and Coeur d’acier A. 2009. Fine-scale cospeciation between *<*i*>*Brachycaudus*<*/i*>* and *<*i*>*Buchnera aphidicola*<*/i*>*: bacterial genome helps define species and evolutionary relationships in aphids. Proceedings. Biological sciences / The Royal Society, 276(1654):187–196.

Karp PD, Paley S, and Romero P. 2002. The Pathway Tools software. Bioinformatics, 18 Suppl 1:S225–232.

Langmead B and Salzberg SL. 2012. Fast gapped-read alignment with Bowtie 2. Nat. Methods, 9(4):357–359.

Latorre A and Manzano-Marín A. 2016. Dissecting genome reduction and trait loss in insect endosymbionts. Annals of the New York Academy of Sciences, pages 1–23.

Lee W, Park J, Lee GS, Lee S, and Akimoto Si. 2013. Taxonomic status of the *Bemisia tabaci* complex (Hemiptera: Aleyrodidae) and reassessment of the number of its constituent species. PloS one, 8(5):e63817.

López-Madrigal S, Latorre A, Moya A, and Gil R. 2015. The link between independent acquisition of intracellular gamma-endosymbionts and concerted evolution in *Tremblaya princeps*. Front. Microbiol., 6(JUN):1–10.

Luan J, Sun X, Fei Z, and Douglas AE. 2018. Maternal inheritance of a single somatic animal cell displayed by the bacteriocyte in the whitefly *Bemisia tabaci*. Curr. Biol., 28(3):459–465.e3.

Luan JB, et al. 2016. Cellular and molecular remodelling of a host cell for vertical transmission of bacterial symbionts. Proc. Biol. Sci., 283(1833):218–230.

Łukasik P, et al. 2018. Multiple origins of interdependent endosymbiotic complexes in a genus of cicadas. Proc. Natl. Acad. Sci., 115(2):E226–E235.

Lynch M. 2010. Evolution of the mutation rate. Trends Genet., 26(8):345–352.

Manzari S and Quicke DLJ. 2006. A cladistic analysis of whiteflies, subfamily Aleyrodinae (Hemiptera: Sternorrhyncha: Aleyrodidae). Journal of Natural History, 40(44-46):2423–2554.

Mao M, Yang X, and Bennett GM. 2018. Evolution of host support for two ancient bacterial symbionts with differentially degraded genomes in a leafhopper host. Proc. Natl. Acad. Sci. U. S. A., 115(50):E11691–E11700.

Marti TM, Kunz C, and Fleck O. 2002. DNA mismatch repair and mutation avoidance pathways. J. Cell. Physiol., 191(1):28–41.

Martin J, Mifsud D, and Rapisarda C. 2000. The whiteflies (Hemiptera: Aleyrodidae) of Europe and the Mediterranean Basin. Bull. Entomol. Res., 90(05).

Martin JH. 1996. Neotropical whiteflies of the subfamily aleurodicinae established in the western palaearctic (Homoptera: Aleyrodidae). J Nat Hist, 30(12):1849–1859.

Martin JH. 2007. Giant whiteflies (Sternorrhyncha, Aleyrodidae):a discussion of their taxonomic and evolutionary significance, with the description of a new species of *Udamoselis* Enderlein from Ecuador. Tijdschr. voor Entomol., 150(1):13–29.

Martin JH. 2008. A revision of *Aleurodicus* Douglas (Sternorrhyncha, Aleyrodidae), with two new genera proposed for palaeotropical natives and an identification guide to world genera of Aleurodicinae. Zootaxa, 1835(1):1–100.

Martin JH and Mound LA. 2007. An annotated check list of the world’s whiteflies (Insecta: Hemiptera: Aleyrodidae). Zootaxa, 1492:1–84.

Martinez-Torres D, Buades C, Latorre A, and Moya A. 2001. Molecular systematics of aphids and their primary endosymbionts. Molecular Phylogenetics and Evolution, 20(3):437–449.

Matsuura Y, et al. 2018. Recurrent symbiont recruitment from fungal parasites in cicadas. Proc. Natl. Acad. Sci., 115(26):E5970–E5979.

Meseguer AS, Coeur d’acier A, Genson G, and Jousselin E. 2015. Unravelling the historical biogeography and diversification dynamics of a highly diverse conifer-feeding aphid genus. Journal of Biogeography, 88(8):1482–1492.

Meseguer AS, Manzano-Marín A, Coeur d’Acier A, Clamens AL, Godefroid M, and Jousselin E. 2017. *Buchnera* has changed flatmate but the repeated replacement of co-obligate symbionts is not associated with the ecological expansions of their aphid hosts. Molecular Ecology, 26(8):2363–2378. 086223.

Misof B, Liu S, Meusemann K, and Peters R. 2014. Phylogenomics resolves the timing and pattern of insect evolution. Science, 346(6210):763–768.

Moran NA. 1996. Accelerated evolution and Muller’s rachet in endosymbiotic bacteria. Proc. Natl. Acad. Sci. U. S. A., 93(7):2873–2878.

Moran NA and Bennett GM. 2014. The tiniest tiny genomes. Annu Rev Microbiol, 68(1):195–215.

Morris JJ, Lenski RE, and Zinser ER. 2012. The Black Queen Hypothesis: evolution of dependencies through adaptive gene loss. MBio, 3(2):e00036–12.

Nabhan AR and Sarkar IN. 2012. The impact of taxon sampling on phylogenetic inference: A review of two decades of controversy. Brief. Bioinform., 13(1):122–134.

Nishimura I, Kurokawa M, Liu L, and Ying BW. 2017. Coordinated changes in mutation and growth rates induced by genome reduction. MBio, 8(4):1–10.

Notredame C, Higgins DG, and Heringa J. 2000. T-Coffee: A novel method for fast and accurate multiple sequence alignment. J. Mol. Biol., 302(1):205–17. URL http://www.ncbi.nlm.nih.gov/pubmed/10964570.

Nováková E, Hypša V, Klein J, Foottit RG, von Dohlen CD, and Moran Na. 2013. Reconstructing the phylogeny of aphids (Hemiptera: Aphididae) using DNA of the obligate symbiont *Buchnera aphidicola*. Molecular Phylogenetics and Evolution, 68(1):42–54.

Okonechnikov K, Golosova O, and Fursov M. 2012. Unipro UGENE: a unified bioinformatics toolkit. Bioinformatics, 28(8):1166–1167.

Ovalle TM, Parsa S, Hernández MP, and Becerra Lopez-Lavalle LA. 2014. Reliable molecular identification of nine tropical whitefly species. Ecol. Evol., 4(19):3778–3787.

Paradis E and Schliep K. 2019. ape 5.0: an environment for modern phylogenetics and evolutionary analyses in R. Bioinformatics, 35(3):526–528.

Patiño-Navarrete R, Moya A, Latorre A, and Peretó J. 2013. Comparative genomics of *Blattabacterium cuenoti* : the frozen legacy of an ancient endosymbiont genome. Genome Biol. Evol., 5(2):351–61.

Quaintance A. 1909. A new genus of Aleyrodidae, with remarks on *Aleyrodes nubifera* Berger and *Aleyrodes citri* Riley & Howard. Tech. Ser. US Dep. Agric. Bur. Entomol., 12:169–174.

R Core Team. 2018. R: A Language and Environment for Statistical Computing. R Foundation for Statistical Computing, Vienna, Austria.

Ranwez V, Douzery EJP, Cambon C, Chantret N, and Delsuc F. 2018. MACSE v2: toolkit for the alignment of coding sequences accounting for frameshifts and stop codons. Mol Biol Evol, 35(10):2582–2584.

Rocha EPC. 2003. An appraisal of the potential for illegitimate recombination in bacterial genomes and its consequences: from duplications to genome reduction. Genome Res., 13(6A):1123–1132.

Rutherford K, Parkhill J, Crook J, Horsnell T, Rice P, Rajandream MA, and Barrell B. 2000. Artemis: sequence visualization and annotation. Bioinformatics, 16(10):944–945.

Santos-Garcia D, et al. 2014. Small but powerful, the primary endosymbiont of moss bugs, *Candidatus* Evansia muelleri, holds a reduced genome with large biosynthetic capabilities. Genome Biol. Evol., 6(7):1875–1893.

Santos-Garcia D, Vargas-Chavez C, Moya A, Latorre A, and Silva FJ. 2015. Genome evolution in the primary endosymbiont of whiteflies sheds light on their divergence. Genome biology and evolution, 7(3):873–88.

Seemann T. 2014. Prokka: rapid prokaryotic genome annotation. Bioinformatics, 30(14):2068–2069.

Shcherbakov DE. 2000. The most primitive whiteflies (Hemiptera; Aleyrodidae; Bernaeinae subfam. nov.) from the Mesozoic of Asia and Burmese amber, with an overview of Burmese amber hemipterans. Bull. nat. Hist. Mus. Lond. (Geol.), 56(June):29–37.

Shereda RD, Kozlov AG, Lohman TM, Cox MM, and Keck JL. 2008. SSB as an organizer/mobilizer of genome maintenance complexes. Crit. Rev. Biochem. Mol. Biol., 43(5):289–318.

Silva FJ and Santos-Garcia D. 2015. Slow and fast evolving endosymbiont lineages: positive correlation between the rates of synonymous and non-synonymous substitution. Front. Microbiol., 6(November):1279.

Sloan DB and Moran NA. 2013. The evolution of genomic instability in the obligate endosymbionts of whiteflies. Genome Biology and Evolution, 5(5):783–793.

Song N, Liang AP, and Bu CP. 2012. A molecular phylogeny of Hemiptera inferred from mitochondrial genome sequences. PloS one, 7(11):e48778.

Szklarzewicz T and Moskal A. 2001. Ultrastructure, distribution, and transmission of endosymbionts in the whitefly *Aleurochiton aceris* Modeer (Insecta, Hemiptera, Aleyrodinea). Protoplasma, 218(1-2):45–53.

Thao ML and Baumann P. 2004a. Evidence for multiple acquisition of *Arsenophonus* by whitefly species (Sternorrhyncha: Aleyrodidae). Current microbiology, 48(2):140–4.

Thao ML and Baumann P. 2004b. Evolutionary relationships of primary prokaryotic endosymbionts of whiteflies and their hosts. Appl Environ Microbiol, 70(6):3401–3406.

Thao ML, Baumann L, and Baumann P. 2004. Organization of the mitochondrial genomes of whiteflies, aphids, and psyllids (Hemiptera, Sternorrhyncha). BMC Evol. Biol., 4:25.

Toft C and Andersson SGE. 2010. Evolutionary microbial genomics: insights into bacterial host adaptation. Nat. Rev. Genet., 11(7):465–475.

Tremblay E. 1959. Osservazioni sulla simbiosi endocellulare dialcuni Aleyrodidae (*Bemisia tabaci* Gennad., *Aleurolobus olivinus* Silv., *Trialeurodes vaporariorum* West.). Bollet. Lab. Entomol. Agrar. Filippo Silvestri Portici, 17:210–246.

Van Leuven JT, Meister RC, Simon C, and McCutcheon JP. 2014. Sympatric speciation in a bacterial endosymbiont results in two genomes with the functionality of one. Cell, 158(6):1270–1280.

Walker BJ, et al. 2014. Pilon: An integrated tool for comprehensive microbial variant detection and genome assembly improvement. PLoS One, 9(11):e112963.

Wang HL, Yang J, Boykin LM, Zhao QY, Wang YJ, Liu SS, and Wang XW. 2014. Developing conversed microsatellite markers and their implications in evolutionary analysis of the *Bemisia tabaci* complex. Scientific reports, 4(SEPTEMBER):6351.

Wernegreen JJ. 2015. Endosymbiont evolution: predictions from theory and surprises from genomes. Ann. N. Y. Acad. Sci., 1360(1):16–35.

Xia X. 2018. DAMBE7: New and Improved Tools for Data Analysis in Molecular Biology and Evolution. Mol. Biol. Evol., 35(6):1550–1552.

Xu XR, Li NN, Bao XY, Douglas AE, and Luan JB. 2019. Patterns of host cell inheritance in the bacterial symbiosis of whiteflies. Insect Sci., pages 1–9.

Yang Z. 2007. PAML 4: Phylogenetic analysis by Maximum Likelihood. Mol Biol Evol, 24(8):1586–1591.

