## Supplementary Material and Methods, and Figures for "*Portiera* gives new clues on the evolutionary history of whiteflies"

#### 1 Supplementary Material and Methods

##### 1.1 Transcriptome assembly and data retrieval

*Dialeurodes citri* (SRR2856996) and *B. tabaci* SSA1 (SRR5109958) transcriptomes were downloaded from the SRA. BLASTN searches against *D. citri* transcriptome were used to retrieve *Portiera*'s genes of interest. *B. tabaci* SSA1 reads were quality filtered with Trimmomatic v0.33 (TruSeq2-PE.fa:2:30:10 LEADING:3 TRAILING:3 SLIDINGWINDOW:4:25 MINLEN:98) (Bolger *et al.*, 2014). Surviving paired-end reads were classified using Kraken v2.0.6-beta with a custom database (Wood and Salzberg, 2014). Kraken 2 custom database included several RefSeq databases (archaea, bacteria, viral, fungi, and protozoa), all sequenced endosymbionts from whiteflies, the genomes of *B. tabaci* MEAM1 and *Acyrthosiphon pisum*, and all complete mitogenomes from whiteflies. Reads classified as *Portiera* were selected and assembled with SPAdes v3.13.0 in RNASeq mode using default parameters (Bankevich *et al.*, 2012; Bushmanova *et al.*, 2019). Resulting *Portiera* transcripts larger than 1 Kb were de-replicated with CD-HIT (EST mode, 96% minimum identity) (Fu *et al.*, 2012), and annotated with Prokka v1.14.5 (–metagenome –gram neg) (Seemann, 2014) to recover the five target genes.
Annotated *Portiera* genomes of *B. tabaci* MEAM1 (NC_018677.1), MED-Q2 (NC_018676.1) (same sequencing procedures as MEAM1), and Asia II 3 (NZ_CP016327.1) species, *Trialeurodes vaporariorum* (LN649236.1), *Aleurodicus dispersus* (LN649255.1), and *A. floccissimus* (LN734649.1) were downloaded from the RefSeq/GenBank and target genes were extracted from each genome.

##### 1.2 *Portiera* lineages Divergence Dating

BEAST v2.5.2 was used to infer a Bayesian posterior consensus tree and the divergence time of the different nodes (Bouckaert *et al.*, 2014). Evolutionary models for each of the pruned alignments were calculated with IQTREE (-m

TESTONLY) (Nguyen *et al.*, 2015). The best model for each alignment was selected based on the Bayesian Information Criterion. BEAUti was used to prepare the requited xml files using the pruned alignments and the obtained evolutionary models. Datasets were partitioned by gene. A lognormal relaxed clock with a Yule speciation process was selected as a prior based on previous a work (Santos-Garcia *et al.*, 2015). The divergence between the subfamilies Aleyrodinae and Aleurodicinae (135?125 Ma) (Drohojowska and Szwedo, 2015) was used as a calibration point using a uniform distribution. Both datasets were first run under the priors to check their possible impact on the estimated dates. Then, four independent runs were conducted for 500 million generations, sampling every 50,000 generations and allowing a pre-burn-in of 1,000 generations. Convergence of the different runs was checked with Tracer v1.6, ensuring that at least an ESS larger than 200 was accomplish for each parameter estimated in each run. Log and Tree files were combined and trimmed with LogCombiner v2.5.2. Combined Tree files were used as input for TreeAnnotator v2.5.2 to obtain the topologies of the trees and their associated parameters, including divergence times and their confidence intervals. Trees were plotted with FigTree v1.4.3 (`https://github.com/rambaut/figtree`).

#### 1.3 *Portiera* lineages Molecular Evolution

Synonymous ($dS$) and non-synonymous ($dN$) substitution ratios and omega ($\omega$) values were calculated in Codeml from PAML v4.7 package (Yang, 2007). To avoid redundancy, only one representative of the *B. tabaci* lineage, the MEAM1 species, was included in the analysis. After quality filtering, 232 of the 235 identified single-copy core OCPs were codon-aligned with MACSE v2.03 as described above and pruned with Gblocks v0.91b (-t=c -b5=n, no gaps allowed). Three branch models available in Codeml were computed: M0 (one $\omega$), M1 (free $\omega$ ratios in each branch) and M2 (two $\omega$, setting *Portiera* from *B. tabaci* and *S. simplex* as the foreground clade). All models were run using a constrained tree option that utilized the species tree obtained with OrthoFinder v2.3.3. Each model was run three times, keeping only the run with the largest likelihood. Likelihood ratio tests (LTR) were used to select the best fitting model, first by comparing model M1 against M2, and later against model M0. Bonferroni correction (two tests, p-value <= 0.025) was used for multiple testing. The pipeline described was implemented in a custom python script.
Divergence times were obtained from the tree that was based on five *Portiera* genes (described above, Figure **??**). These estimates were used to calculate the number of nucleotide substitutions per site per year ($dS/t$ and $dN/t$) in each whitefly lineage. Before any statistical test, OCPs with $dS$, $dN$ or $\omega$ values below percentile 1 or above percentile 99 were discarded. Also, OCPs with $omega$ values greater than 10 were discarded. Raw and log-transformed data were tested for normality and homoscedasticity (Shapiro's and Levene's test, respectively). As $dS/t$, $dN/t$, and $\omega$ values were not normally distributed or homoscedastic, the non-parametric Kruskal-Wallis rank-sum and post-hoc pairwise Wilcoxon rank sum tests (with Benjamini & Hochberg correction) were used. All statistical analyses were performed in R.

#### 1.4 Repeats and intergenic regions analysis

Representatives genomes of insects' endosymbionts with reported large intergenic regions, genome instability, or lacking *dnaQ* were downloaded: *Ca.* Tremblaya princeps (NC_01729) - the primary endosymbiont of the citrus mealybug *Planococcus citri*, *Ca.* Uzinura diaspidicola (NC_020135) ? the primary endosymbiont of scale insects from the family Diaspididae, and *Ca.* Hodgkinia cicadicola TETUND1 and 2 (CP007232, CP007233) - bacterial symbionts of cicada species of the genus *Tettigades*. We also included *Portiera* genomes from *B. tabaci* MEAM1 (NC_018677.1) and MED (NC_018676.1), and *T. vaporariorum* (LN649236.1). For standardization purposes, UGENE v1.28.1 was used to predict not only direct and inverted repeats (ugene find-repeats –identity=95 –inverted=[true—false] –max-distance=500000 –min-length=50 –thread-count=12 –exclude-tandems=true), but also tandem repeats (UGENE GUI Find Tandems wit Tandem presets: Mini-satellite, length 7-30 nt, minimum tandem size 9 nt, minimum repeat count 3) (Okonechnikov *et al.*, 2012). Intergenic regions were extracted with a custom python script and their length and GC content were computed with infoseq from EMBOSS v6.6.0.0 (Rice *et al.*, 2000). As length distributions were found not to be normal and homoscedastic, non-parametric tests were used as described above) Statistical analyses were performed in R.

#### 1.5 Mitochondrion assembly, annotation, and molecular evolution

For accurate assembly of the mitochondrial genomes of *S. simplex* and *P. mori*, reads classified as "Insect" by Kraken2 were extracted and assembled with SPAdes v3.13.0 (–sc –careful). *mtCOI*s fragments obtained by PCR screening were used as a query for a BLASTN search against the assembly. Two contigs larger than 15Kb containing a full *mtCOI* gene were recovered. The first contig presented a full *mtCOI* gene with a nucleotide identity higher than 97% to *P. mori* (LR739216). The second contig harboring a full *mtCOI* gene had *Acaudaleyrodes rachipora* (LR739211) and *B. reyesi* (LR739074) as the two best hits. Further confirmation was conducted with the BOLD Identification System (Ratnasingham and Hebert, 2013). The first *mtCOI* full gene was classified again as *P. mori* (98.97% similarity to BOLD record GMESH030-14). The second *mtCOI* full gene was classified as *S. simplex* (100% similarity to BOLD record GBMHH7908-15). Both contigs were circularized with *Gap5* and corrected with Pilon v1.23 (Walker *et al.*, 2014) as described above. Mitochondrial genomes annotation was performed with MITOS v2 web server (`http://mitos2.bioinf.uni-leipzig.de`) setting the genetic code to Invertebrate. Annotations were downloaded and manually revised in Artemis v16 (Rutherford *et al.*, 2000).

*A. dispersus* (KR063274), *T. vaporariorum* (NC_006280), and *B. tabaci* MED (MH205752) mitogenomes were downloaded from NCBI. Full genes were extracted from the downloaded and the newly assembled *P. mori* and *S. simplex* mitogenomes. Due to annotation differences, only 12 mitochondrial genes were shared between the different genomes. Extracted genes were codon-aligned with MACSE v2.03 (genetic code set up to 5) and pruned with Gblocks v0.91b. Saturation of the phylogenetic signal was assessed with DAMBE v7.2.3 as described above. Finally, pruned alignments were used as an input for Codeml to calculate $dS$, $dN$, and $\omega$ values. Since only one *Aleurodicus* species was included, the divergence of this lineage was set up to 129.35 Myr (the estimated time for the split between the Aleurodicinae and the Aleyrodinae families). Divergence times of the other four lineages were the same as described above. Statistical analyses and data cleaning were conducted as described above but this time the data were normally distributed and one-way ANOVA with Tukey's post-hoc tests were applied when required.

#### 2 Supplementary Figures

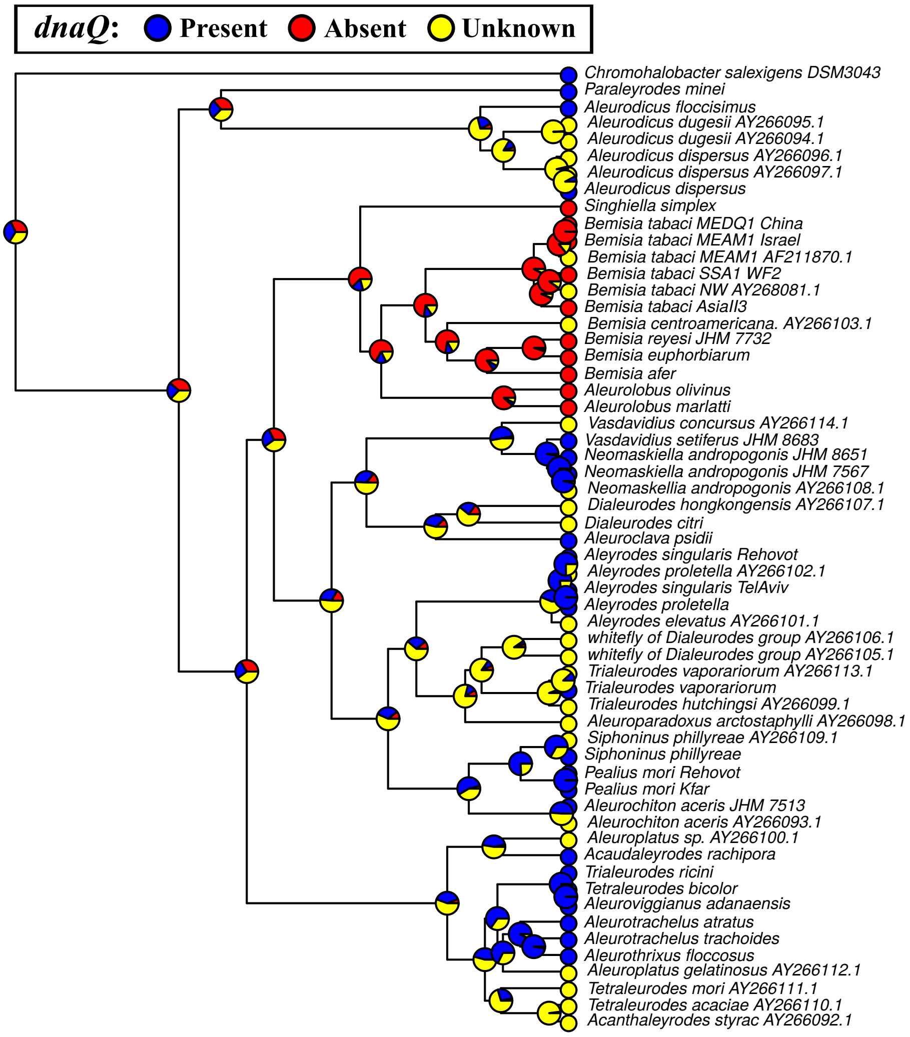


Figure S1: DNA polymerase III proofreading subunit (*dnaQ*) presence or absence using the two *Portiera* genes-based tree (the *16S* and *23S* rRNA genes). The ancestral state for each node was estimated using the ace function form the ape package (Paradis and Schliep, 2019) in R (R Core Team, 2018). Pie charts at the nodes represent the posterior probability for the presence (blue), absence (red), or unknown state (yellow). Filled circles at the tips represent *dnaQ* successful amplifications (blue), no amplification (red), or no data/untested (yellow).

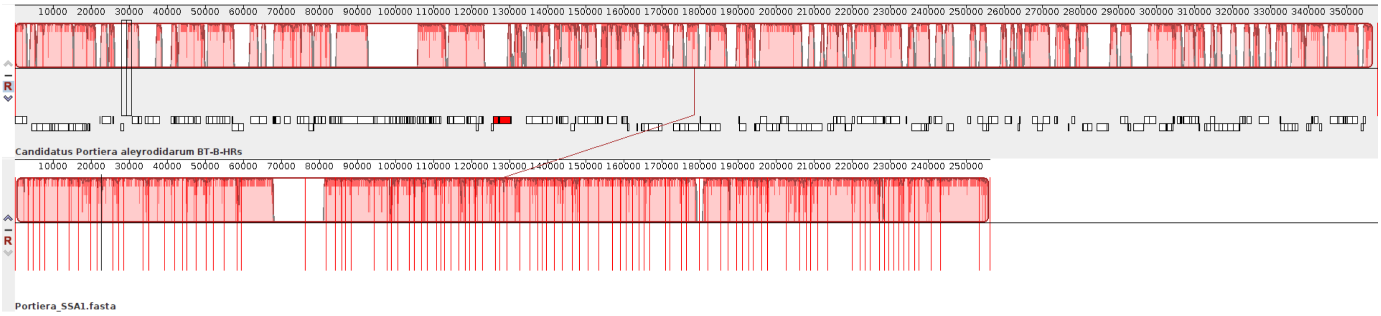


Figure S2: Syntenyc blocks shared by *Bemisia tabaci* MEAM1 species genome and *B. tabaci* SSA1 species recovered transcripts. Syntenic blocks were computed using the progressive Mauve aligner snapshot 2015-02-13 (Darling *et al.*, 2011).

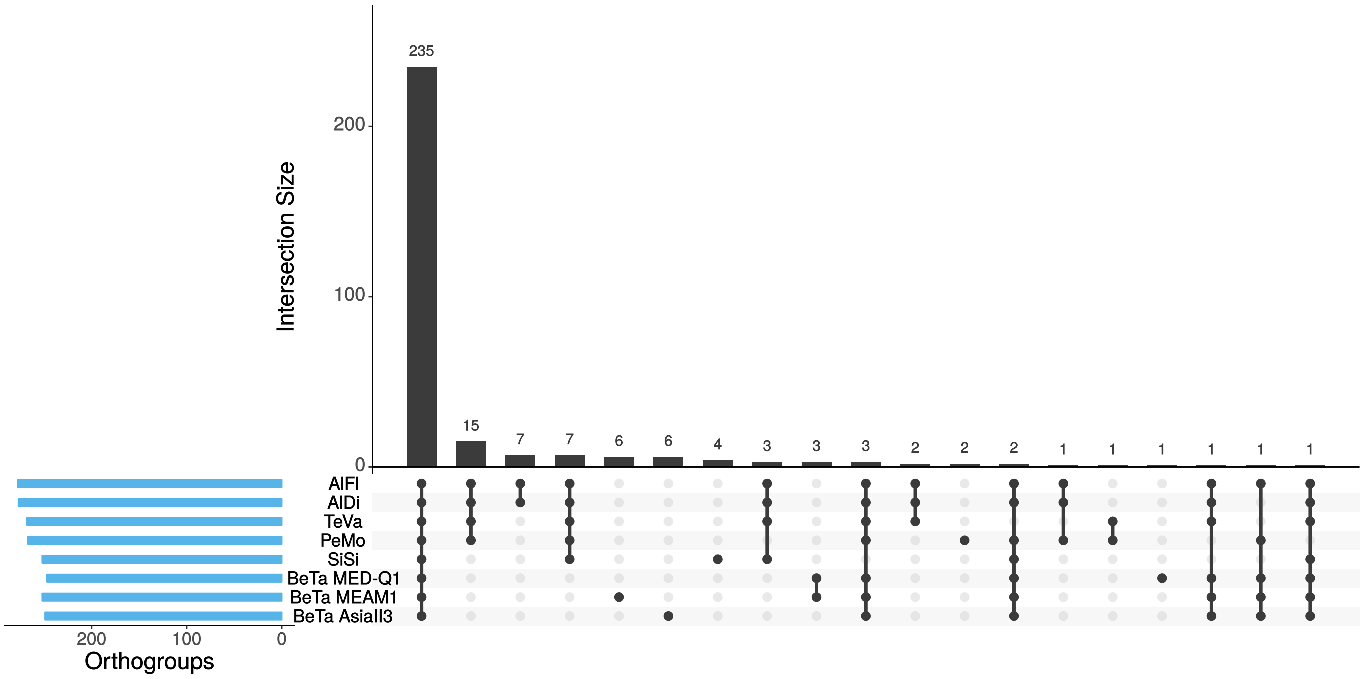


Figure S3: Upset plot showing the core, shared, and specific Orthologous Clusters of Proteins (OCPs) between *Portiera* from *Aleurodicus floccissimus* (AlFi), *A. dispersus* (AlDi), *Trialeurodes vaporariorum* (TeVa), *Pealius mori* (PeMo), *Singhiella simplex* (SiSi), and *Bemisia tabaci* (BeTa) MED-Q1, MEAM1, and AsiaII3 species.

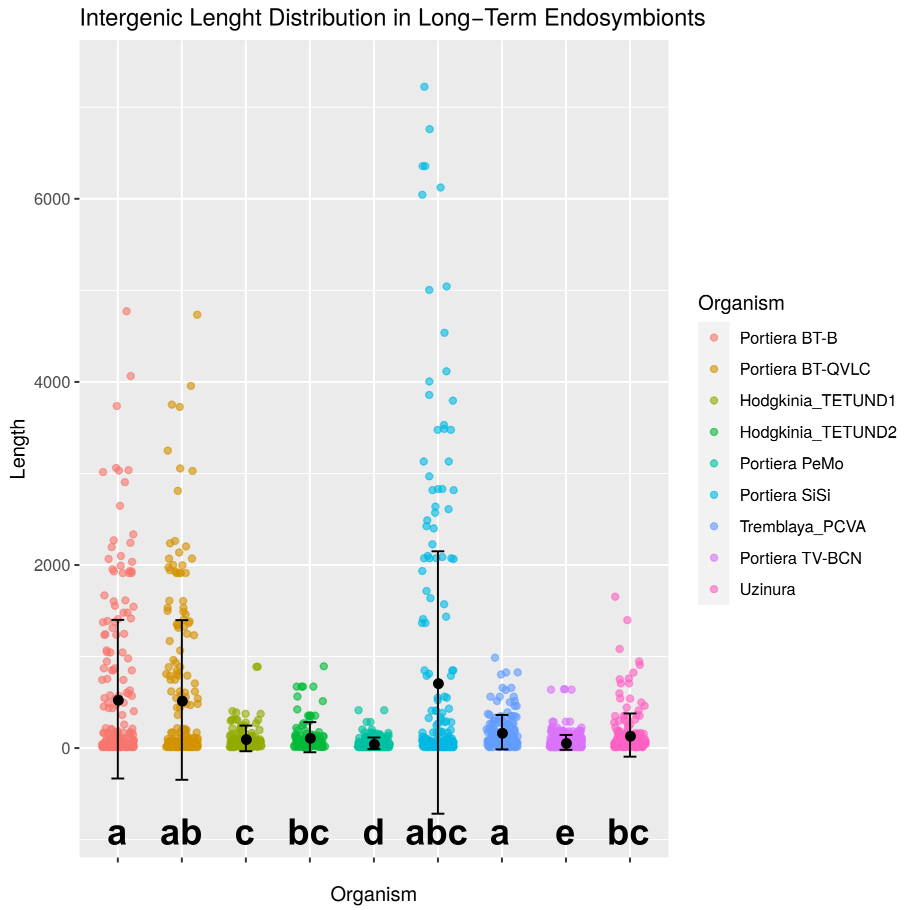


Figure S4: Length distribution of the intergenic regions from selected *Portiera aleyrodidarum*, *Uzinura diaspidicola*, *Tremblaya princeps*, and *Hodgkinia cicadicola* genomes. Different letters indicate significant statistical differences between intergenic regions length distributions (non-parametric Kruskal-Wallis and Wilcoxon post-hoc pairwise tests).

### References

Bankevich A, *et al.* 2012. SPAdes: a new genome assembly algorithm and its applications to single-cell sequencing. *J. Comput. Biol.*, **19**(5):455–77.

Bolger AM, Lohse M, and Usadel B. 2014. Trimmomatic: a flexible trimmer for Illumina sequence data. *Bioinformatics*, **30**(15):2114–2120.

Bouckaert R, *et al.* 2014. BEAST 2: A software platform for Bayesian evolutionary analysis. *PLoS Comput Biol*, **10**(4):e1003537.

Bushmanova E, Antipov D, Lapidus A, and Przhibelskiy AD. 2019. rnaspades: a de novo transcriptome assembler and its application to rna-seq data. *GigaScience*, **8**(9).

Darling AE, Tritt A, Eisen Ja, and Facciotti MT. 2011. Mauve assembly metrics. *Bioinformatics*, **27**(19):2756–2757.

Drohojowska J and Szwedo J. 2015. Early Cretaceous Aleyrodidae (Hemiptera: Sternorrhyncha) from the Lebanese amber. *Cretac. Res.*, **52**:368–389.

Fu L, Niu B, Zhu Z, Wu S, and Li W. 2012. CD-HIT: accelerated for clustering the next-generation sequencing data. *Bioinformatics*, **28**(23):3150–2.

Nguyen LT, Schmidt HA, von Haeseler A, and Minh BQ. 2015. IQ-TREE: A fast and effective stochastic algorithm for estimating Maximum-Likelihood phylogenies. *Mol Biol Evol*, **32**(1):268–274.

Okonechnikov K, Golosova O, and Fursov M. 2012. Unipro UGENE: a unified bioinformatics toolkit. *Bioinformatics*, **28**(8):1166–1167.

Paradis E and Schliep K. 2019. ape 5.0: an environment for modern phylogenetics and evolutionary analyses in R. *Bioinformatics*, **35**(3):526–528.

R Core Team. 2018. *R: A Language and Environment for Statistical Computing.* R Foundation for Statistical Computing, Vienna, Austria.

Ratnasingham S and Hebert PDN. 2013. A DNA-based registry for all animal species: The Barcode Index Number (BIN) system. *PLoS One*, **8**(7):e66213.

Rice P, Longden I, and Bleasby A. 2000. EMBOSS: The European Molecular Biology Open Software Suite. *Trends Genet*, **16**(1):276–277.

Rutherford K, Parkhill J, Crook J, Horsnell T, Rice P, Rajandream MA, and Barrell B. 2000. Artemis: sequence visualization and annotation. *Bioinformatics*, **16**(10):944–945.

Santos-Garcia D, Vargas-Chavez C, Moya A, Latorre A, and Silva FJ. 2015. Genome evolution in the primary endosymbiont of whiteflies sheds light on their divergence. *Genome biology and evolution*, **7**(3):873–88.

Seemann T. 2014. Prokka: rapid prokaryotic genome annotation. *Bioinformatics*, **30**(14):2068–2069.

Walker BJ, *et al.* 2014. Pilon: An integrated tool for comprehensive microbial variant detection and genome assembly improvement. *PLoS One*, **9**(11):e112963.

Wood DE and Salzberg SL. 2014. Kraken: Ultrafast metagenomic sequence classification using exact alignments. *Genome Biology*, **15**(3). `/www.pubmedcentral.nih.gov/articlerender.fcgi?artid=3006164{&}tool=pmcentrez{&}rendertype=abstract.`

Yang Z. 2007. PAML 4: Phylogenetic analysis by Maximum Likelihood. *Mol Biol Evol*, **24**(8):1586–1591.
